# A widespread gut bacterial lineage distinguished by redox metabolism and phage defense

**DOI:** 10.64898/2026.03.31.715625

**Authors:** Cecilia Noecker, Lu Guo, Cyrille Daté, Nitee Rai, Fadima Daramy, Luis A. Ramirez Hernandez, Than S. Kyaw, Kai R. Trepka, Chhedi Lal Gupta, Connie W.Y. Ha, Joel Babdor, Matthew H. Spitzer, Peter J. Turnbaugh

## Abstract

Genomic variation within gut microbial species can have consequences for host health and disease. However, for low abundance species, these variations can be difficult to capture by both culture-dependent and -independent approaches. Here, we focus on the prevalent but low abundance gut Actinomycetota *Eggerthella lenta*. We developed a selective media for sensitive and specific isolation of *E. lenta* from human stool. Genomes from 87 new *E. lenta* isolates were combined with prior high-quality assemblies, shedding light on within-species functional diversity. Phylogenetic analysis revealed a broadly distributed subclade, which we refer to as *E. lenta* Group B. This lineage was differentiated by its metabolic potential and bacteriophage defense, though mobile elements were shared broadly across the species. Notably, Group B was positively associated with intestinal inflammation in subjects with inflammatory bowel disease. Overall, these results emphasize the importance of bacterial population structure in host-microbiome interactions and provide a framework to study low-abundance gut taxa.

**HIGHLIGHTS:**

- Selective media enables *E. lenta* isolation and reveals high prevalence in humans
- Discovery of a distinctive lineage within *E. lenta* undergoing genome reduction
- *E. lenta* Group B has altered metabolism, phage defense, and disease associations
- A widespread conjugative plasmid could enable improved genetics

## INTRODUCTION

In the human gut microbiota, members of the same microbial species may differ widely in their functional capabilities. In a classic example, different strains of *Escherichia coli* may act as commensals, probiotics, or pathogens depending on carriage of specific accessory genes^1,2^. Similarly, within-species differences in other gut taxa can impact immune regulation, drug metabolism, dietary response, ecological interactions, and other aspects of community function^3–6^. Correspondingly, the population structure of gut bacterial strains has been linked with the occurrence and/or progression of inflammatory bowel disease^7,8^ and colorectal cancer^9^. Such population structure may develop via a variety of mutational and selection processes on both short and long time scales, incorporating nucleotide mutation, recombination, and multiple types of horizontal gene transfer^10–12^.

Much of the current scientific understanding of the within-species population structure of gut microorganisms comes from metagenomic sequencing, with a focus on the most abundant taxonomic groups. Metagenomic assembly across thousands of samples revealed gut microbial strain variation across gradients of diet, geography, and lifestyle^13–15^, and that strain variation can be driven by bacteriophage interactions, interbacterial antagonism^16–18^, and differences in transmissibility^19,20^, among other factors. However, since most reads in a metagenomic study come from the dominant Bacteroidota and Bacillota phyla, tracking strain populations in taxa that form a smaller fraction of the community presents technical challenges unless extremely deep sequencing has been performed.

One such low-abundance gut microorganism is *Eggerthella lenta,* a prevalent member of the Actinomyctetota phylum^21^ that is likely further underrepresented in metagenomic databases due to its lysis resistance and high GC content^22^. Even so, *E. lenta* has consequential links with human health: its abundance is elevated in a variety of chronic conditions, with a defined role in autoimmunity including inflammatory bowel disease and rheumatoid arthritis^23,24^. *E. lenta* can also directly impact other aspects of human physiology including drug metabolism and bile acid signaling, among others^23–28^. Many of these impacts are mediated by strain-variable gene families^29^, suggesting that different strains of *E. lenta* may have disparate impacts on health. Supporting this view, one recent study was able to define two *E. lenta* lineages in metagenomic data using a small number of core gene read alignments, revealing a lineage associated with lower levels of fecal calprotectin in inflammatory bowel disease patients^7^.

In this study, we sought to accomplish the following key goals: (*i*) develop and validate a selective media to efficiently isolate *E. lenta* from human stool samples; (*ii*) compile an atlas of high-quality *E. lenta* genomes to describe large-scale trends in its genome evolution across populations; and (*iii*) examine factors influencing *E. lenta* population structure and strain diversity. We found a dynamic pangenome, distinctive mobile elements, and a large and broadly distributed lineage emerging within the *E. lenta* species. This clade is distinguished by its genomic properties, metabolic capabilities, and antiviral defense systems, and has distinct associations with intestinal inflammation. These findings provide context for *E. lenta*’s strain-variable impacts in the gut and establish a substantially expanded resource for further studies of its genetics and evolution.

## RESULTS

### Establishing a repository of *E. lenta* genomes

Due to the underrepresentation of *E. lenta* in metagenomic and cultureomics datasets^21,22^, we sought to develop a targeted method to isolate *E. lenta.* Based on our previous metabolic modeling analysis of *E. lenta*^30^, we designed a media (ESM, *Eggerthella* Selective Media) lacking common media components dispensable for *E. lenta* growth (**Table S1**). Growth of 9 *Eggerthellaceae* isolates in monoculture revealed that only the two tested *Eggerthella* strains grew in ESM (**Figure S1A**). Growth of stool-derived bacteria on solid ESM agar followed by 16S rRNA gene sequencing further confirmed that this ESM is specific to *Eggerthella* (98.7±1.1% of reads from total 10^−5^ plate contents, *n*=3 samples; **Figure S1B,C**). We then expanded our efforts to stool samples collected from a cohort of healthy adults in the San Francisco Bay Area^31^, successfully isolating *E. lenta* from 90.1% (82/91) of the samples (**Table S2, Figure S1D**). The prevalence of *E. lenta* by culturing on ESM was slightly higher than estimated by metagenomics, and *E. lenta* abundance estimates were highly correlated between the two methods (**Figures S1E**).

Overall, we built a curated repository of 284 high-quality *E. lenta* genomes (**Table S3, Figure 1A, Figure S2A**). These genomes derived from hybrid short- and long-read sequencing of our new isolates (after 99.9% dereplication, *n*=87 isolates from 74 individuals), 81 previously published isolate genomes, and 116 metagenome-assembled genomes (MAGs) with at least 75% completeness and less than 3% contamination (**Figure S2B,C**). We used the consensus of multiple quality-checking tools to assess completeness and contamination and to remove low-quality genomes (**Methods**). In the final set of genomes, the vast majority were sourced from human subjects (98.9%) and gastrointestinal tract samples (93.3%), supporting the prevailing view of *E. lenta* as a human gut-adapted commensal (**Table S3**).

**Figure 1.**
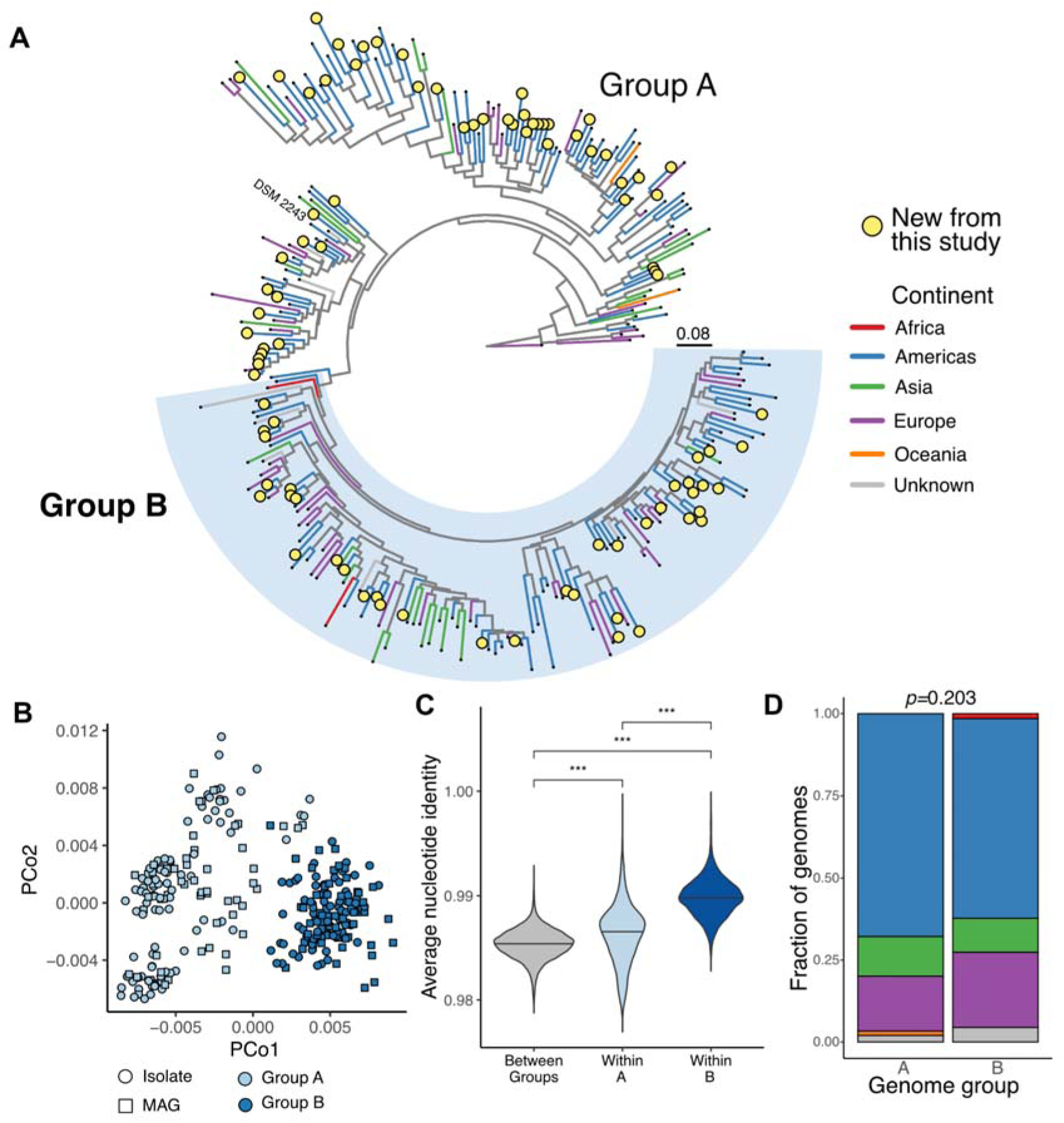
A genome atlas of the human gut-associated species *Eggerthella lenta* reveals subspecies population structure. **(A)** Marker-gene based phylogeny of *E. lenta* genomes analyzed in this study. Newly sequenced isolates are shown as yellow circles. The type strain is labeled as “DSM 2243”. The legend indicates branch length in substitutions per site. The newly defined Group B clade is labeled in light blue (*n=*135 Group B genomes, *n=*149 Group A). **(B)** Principal coordinate analysis plots representing *E. lenta* genomes in terms of overall sequence similarity (Mash distances), colored by assignment of genomes into Group A or B. **(C)** Pairwise comparisons of whole genome average nucleotide identity between genomes within each group and between groups. Similarities were compared using a Wilcoxon rank-sum test. **(D)** Comparison of Groups A and B in terms of their continent of origin, with no significant difference in distribution between the two groups (Fisher’s exact test).

We constructed an *E. lenta* pangenome from this dataset using eggNOG orthologous gene family annotations^32^, finding a total of 5,932 gene families. On average, over a quarter of any strain’s genome consisted of accessory genes (25.9%). Accessory gene families spanned a variety of gene functions and continued to accumulate even after 284 genomes (**Figure S2D-F**), including a large but variable number of genes involved in anaerobic respiration across all genomes (**Figure S2G).** The pangenome was estimated to be open based on Heaps’ law (11=0.90, **Figure S2D-E, Methods**), emphasizing the genomic diversity across the *E. lenta* species. The level of saturation and the functional distribution of the pangenome were comparable when the analysis was restricted to only include isolate genomes (**Figure S2E-F**). Furthermore, a phylogenetic pangenome model constructed with the tool Panstripe (**Methods**) confirmed a high rate of gene gain and loss across most branches of the tree (**Figure S2H**).

### A geographically dispersed lineage within *E. lenta*

Multiple methods identified clear population structure within the *E. lenta* species. Hierarchical clustering based on genome-wide sequence similarity^33^ revealed two main genome clusters (**Figures 1A,B and S3A,B**). However, only one of these clusters was represented by a single monophyletic lineage (**Figure 1A**), referred to hereafter as *E. lenta* Group B. We refer to the other group of genomes as *E. lenta* Group A (“<u>A</u>ll others”), as this cluster contains multiple paraphyletic lineages. Group B was clearly differentiated from the rest of the species under phylogenetic reconstructions using multiple core gene databases (**Figure S3C**, **Methods**), by whole-genome average nucleotide identity (ANI; **Figure S3D,E**) and gene family presence/absence (**Figure S3F**). Notably, this clade was not sufficiently diverged to represent a different species: the smallest observed ANI between any pair of *E. lenta* genomes is 97.6% (**Figure 1C**). Consistent with these results, the 16S rRNA gene was highly conserved across all sequenced genomes (average identity of 99.2% between 16S rRNA gene pairs, minimum of 91.5%).

Surprisingly, the population structure of *E. lenta* was only weakly associated with the geographical location of sampling, as both groups were distributed broadly across 4 continents (**Figures 1A,D**). Geographical source was modestly associated with genome variation across the overall collection by multiple independent metrics (**Figure S3G-I**). This variation was not associated with any specific genome source within the repository (**Figure S3J**). The geographical signal was weaker than the association observed in genomes from another closely related *Eggerthellaceae* species (**Figure S3K**). The continent of birth was also weakly associated with *E. lenta* sequence variation (**Figure S3L**), but did not significantly differ between genome groups (*p=*0.08, Fisher exact test). Thus, geography appears to play a statistically significant but relatively minor role in shaping *E. lenta* strain variation.

### *E. lenta* Group B has a reduced genome and distinctive metabolic accessory genes

Next, we sought to evaluate differences in genome properties and metabolism that could potentially lead to niche segregation between *E. lenta* lineages. We observed that Group B genomes were significantly smaller than other genomes (by 115 kb or 3.4% on average, **Figure 2A**). They also encoded significantly fewer genes per genome (**Figure 2B**) and had a relatively larger proportion of the genome devoted to core genes (**Figure 2C**). Taken together, these results are indicative of genome reduction within *E. lenta* Group B.

**Figure 2.**
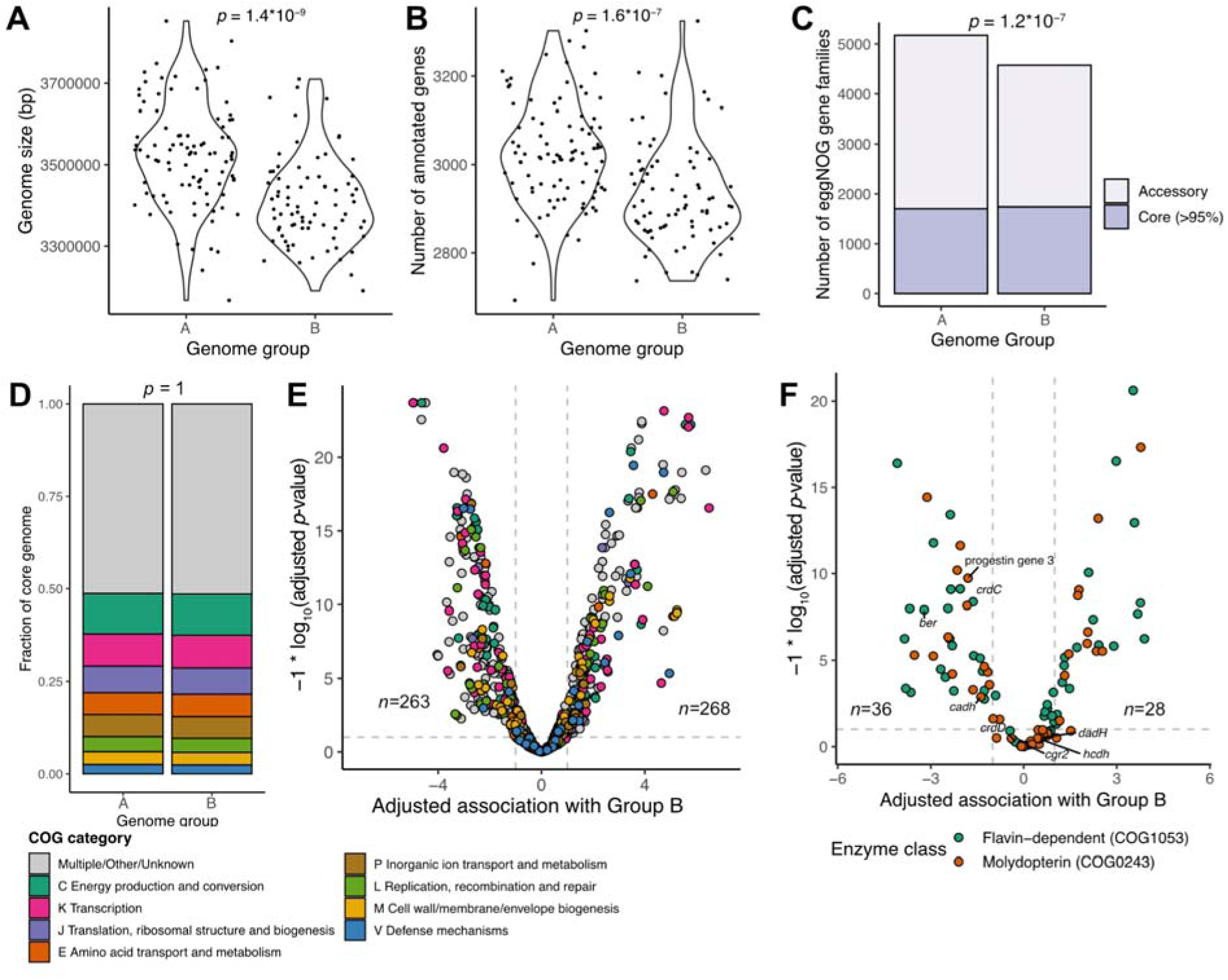
*E. lenta* Group B differs in genomic properties and in specific gene families. **(A-B)** Isolate genomes from *E. lenta* Group B have both a smaller genome size **(A)** and a smaller number of protein-coding genes **(B)** compared with Group A isolate genomes (*n=*135 Group B genomes, *n=*149 Group A). *p-*values are based on Wilcoxon rank-sum tests. **(C)** Comparison of the pangenome of *E. lenta* Groups A and B. The core genome is similar in size between them; however, Group B has a smaller accessory genome despite consisting of a similar number of genomes. The indicated *p-*value is for enrichment of core genes in Group B compared to Group A (Fisher’s exact test). **(D)** Distribution of high-level gene functions in the core genome is similar between Groups A and B (Fisher’s exact test). **(E)** Association of eggNOG gene families with either Group A or B genomes, based on a logistic regression model adjusting for genome size and completeness. 7 gene families with low prevalence and exclusively found in one group are not shown. The number of significantly enriched and depleted gene families is indicated. 522 gene families were not significantly associated. **(F**) Association of anaerobic respiration reductase enzyme genes with either Group A or B genomes, based on a logistic regression model adjusting for genome size and completeness. Genes were identified based on eggNOG annotations and subdivided into higher resolution families using ProteinOrtho. Genes encoding enzymes with known specificity are labeled, including *cgr2* (cardenolide metabolism)^6,21^, *dadh, cadh,* and *hcdh* (catechol metabolism)^26,106^, *ber* (enterolignan metabolism)^34^, *crdC-D* (cinnamic acid metabolism)^36^, and genes involved in metabolism of progestins^35^. One gene cluster only found in Group B (estimate > 19, *p*-value not significant) is not shown. The number of significantly enriched and depleted gene families is indicated. 48 gene families were not significantly associated with Group B.

Genes involved in primary metabolism were largely conserved across the species. The two groups did not differ significantly in the composition of their core genomes across broad functional categories (**Figure 2D**, *p*=1, Fisher exact test). We examined key metabolic genes and pathways identified in our prior metabolic modeling study^30^, including the arginine and agmatine deiminase pathways and pyruvate-flavodoxin oxidoreductase for the use of acetate as a biosynthetic carbon source. All were fully conserved except for a single isolate missing complete pathways for agmatine catabolism (**Table S4**). Auxotrophies for tryptophan and riboflavin were also conserved across all strains. Interestingly, five genes with annotations related to heme and/or cobalamin cofactor biosynthesis were predicted to be essential for *E. lenta* growth in our previous modeling study but were depleted in most Group B genomes, again consistent with reductive genome evolution (**Table S4**).

*E. lenta* Group B also differed from Group A in other areas of metabolism. Overall, 531 out of 1,053 accessory gene families were either enriched (*n=*268) or depleted (*n=*263) in Group B (**Figure 2E**, logistic regression FDR-adjusted *p*<0.1, estimate>2). In particular, the set of respiratory reductase enzymes differed markedly between the two groups. These enzymes, which encompass two major families (flavin-dependent and molybdopterin-dependent), include several that are unique to *E. lenta* and have been linked to host physiology and signaling in various ways^6,23,25,26,34–36^. While all *E. lenta* genomes harbored large repertoires of these enzymes (**Figure S2G**), most (57%) individual enzymes were either enriched or depleted in Group B (**Figure 2F**, 28 out of 112 enriched, 36 depleted, logistic regression FDR-adjusted *p* < 0.1 and estimate > 2). Interestingly, several previously studied reductase genes, including *ber* (encoding an enzyme responsible for lignan metabolism^34^) and genes involved in conversion of glucocorticoids into progestins^35^ are strongly depleted in *E. lenta* Group B (**Figure 2F, Table S4**). Others, however, including *cgr2*^6,21^ (responsible for cardenolide metabolism), are distributed broadly across the species phylogeny. Overall, despite *E. lenta*’s apparent high levels of strain variation across the phylogeny^29^, Group B has consistent functional differences from the rest of the species.

### Two loci linked to bacteriophage defense and redox metabolism distinguish *E. lenta* Group B

Given these broad differences in overall gene content, we next performed a systematic search for genes that are widespread in *E. lenta* Group B but nearly absent in Group A. This pattern would be suggestive of acquisition near the time of divergence under a maximum parsimony model^37^ (**Methods**) and are thus more likely to play a causal role in *E. lenta* genome evolution. This analysis revealed 16 genes strongly linked to Group B, most of which (*n=*11) were clustered in just two genomic loci (**Figure S4**).

One major locus appears to be a defense island (**Figure 3A and S4C**), found in 89.5% of Group B isolate genomes compared with 2.2% Group A (odds ratio=348.2, *p<*10^−15^, Fisher exact test). Genes in the region include multiple with potential roles in phage defense: a putative Type II restriction endonuclease; a cytosine methyltransferase; a large gene (3,039 bp) with domains related to RNA helicase, primase, and endonuclease activity; and a Mrr-like Type IV restriction enzyme. This gene assemblage does not appear to conform with the structure of any well-studied defense system^38^.

**Figure 3.**
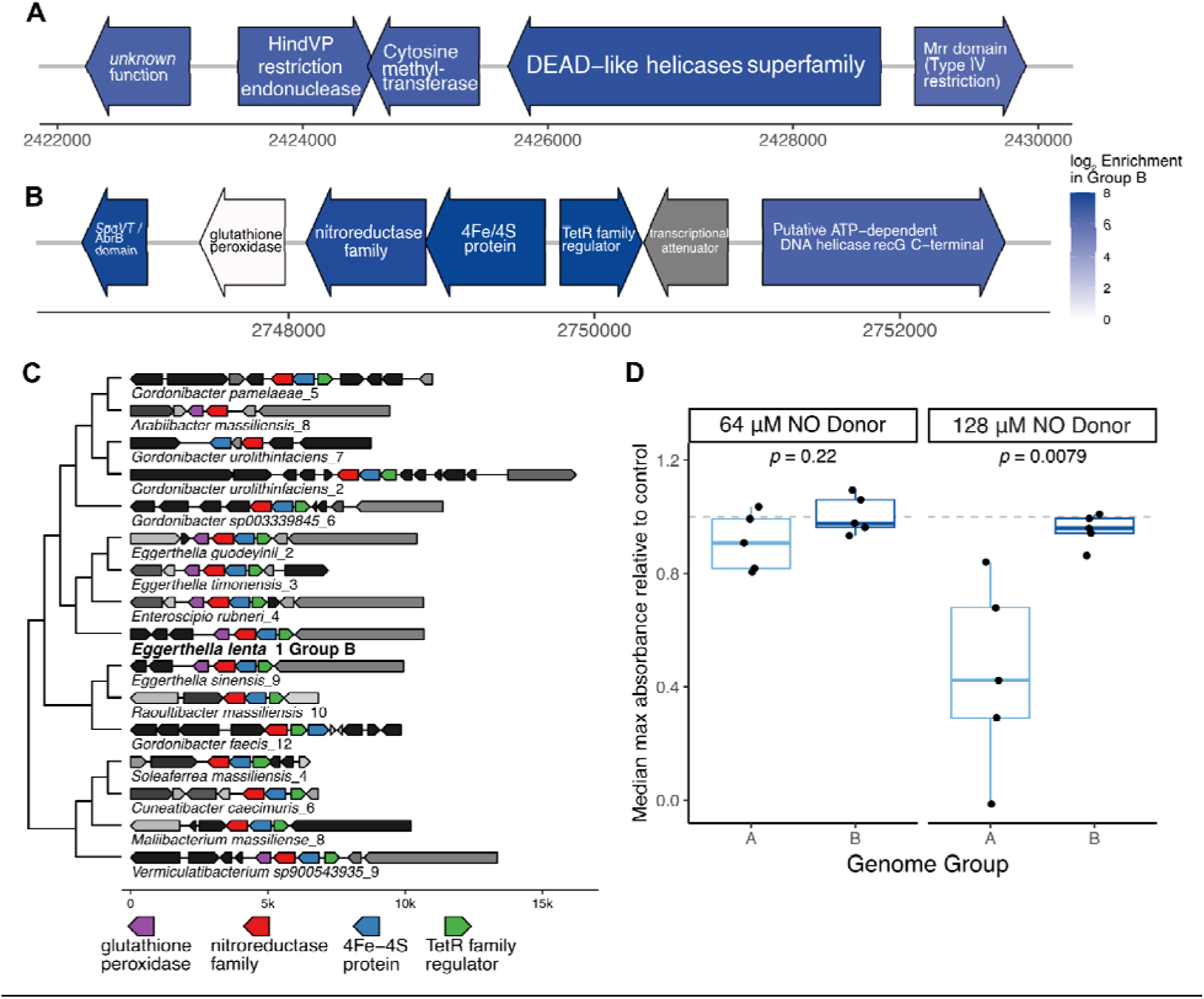
Genomic regions linked to phage defense and redox metabolism are specific to *E. lenta* Group B. **(A,B)** Gene maps of two of the most enriched regions in Group B genomes compared with Group A. The region shown in **(A)** appears to be a defense island. The region shown in **(B)** contains an operon carrying genes annotated as a putative nitroreductase, an iron-sulfur binding enzyme of unknown function, and a putative glutathione peroxidase. Genes are colored by their log enrichment in Group B genomes, based on a logistic regression model accounting for genome size and MAG status. The peroxidase gene in **(B)** does not appear enriched because another copy of this gene family appears elsewhere in the core *E. lenta* genome. Maps of these regions with additional genomic context are included in **Figure S4. (C)** Comparison of the region shown in **(B)** with homologous regions identified in other *Eggerthellaceae* and in representatives of class Clostridia. The region is found widely, though not universally, in other *Eggerthellaceae*, and is also common in Clostridia. Homologous regions were identified and used to construct the associated gene tree using the tool *fast.genomics*. The region found in *E. lenta* Group B is labeled in bold. The phylogenetic tree is based on the nitroreductase gene amino acid sequences. **(D)** Differences in growth inhibition by nitric oxide in Group A and B *E. lenta.* Isolates (*n=*5 from each group) were inoculated into rich media containing initial 64 or 128 µM concentrations of the nitric oxide donor diethylamine NONOate (Wilcoxon rank-sum test). The *y*-axis represents the fraction of the maximum absorbance reached by that strain in control media.

A second region is also found near-universally in Group B and rarely in other genomes **(Figure 3B**, 97.4% Group B isolate prevalence compared with 2.2% Group A, odds ratio=1264.1, *p<*10^−15^, Fisher exact test). It contains 4 out of the top 6 most strongly associated gene families (**Figure S4E**, **Table S4**). Among other genes, this region includes an operon containing genes for two small iron-sulfur binding domain enzymes, one of which is annotated as a nitroreductase family protein, and a glutathione peroxidase. Interestingly, a homologous operon was found in a wide subset of other *Eggerthellaceae* (**Figure 3C**), suggesting that it may have been lost in genus *Eggerthella* but then reacquired by *E. lenta* Group B. More distant homologs are also found in diverse members of class Clostridia (**Figure 3C**).

Based on the presence of a putative nitroreductase gene as well as the adjacent peroxidase, we hypothesized that it could be involved in maintaining anaerobic respiration in the context of free radical nitrosative and/or oxidative stress, as has been observed in some organisms^39,40^, although in others this enzyme class is linked with greater susceptibility to oxidative stress^41^. As an initial test, we evaluated whether *E. lenta* strains vary in their susceptibility to nitrosative stress by quantifying growth inhibition in the presence of a nitric oxide donor (**Methods**). As predicted, *E. lenta* Group B strains that carried this operon displayed less growth inhibition in the presence of nitric oxide than other strains without this operon (**Figure 3D**).

### Defense system repertoires are lineage-specific while a common conjugative plasmid is found across the species

Next, we considered whether processes of genetic isolation and/or horizontal gene transfer could influence *E. lenta* diversification and evolution. 35 gene families in the categories of defense (V) and recombination/repair (L) were differentially prevalent between Group B and the rest of the species (**Figure 2E**). Therefore, we evaluated the distribution of defense systems and mobile elements in our repository in a more targeted way.

We uncovered a diverse array of defense systems in the pangenome using the Hidden Markov Model-based tool DefenseFinder^38^. Most of these were strain-variable (**Figure 4A**): the only system in the core genome was AbiE, an abortive infection toxin-antitoxin system^42^. Group B genomes harbored a slightly higher total number of defense systems (**Figure 4B**), despite their relatively smaller genome size (**Figure 2A**). Nearly all Group B genomes (96%) encoded at least two annotated restriction-modification (RM) systems, significantly more than the 62.0% of genomes in Group A (**Figure 4C**, number of RM systems *p=*0.016, Poisson regression model). They were also more likely to possess Type IV restriction enzymes and less likely to carry Types II and III (**Figure 4D**). Similarly, CRISPR-Cas systems were also nearly universal in Group B (98.7% of isolates), significantly more than in Group A (68.5%; *p=*3×10^−8^, logistic regression model).

**Figure 4.**
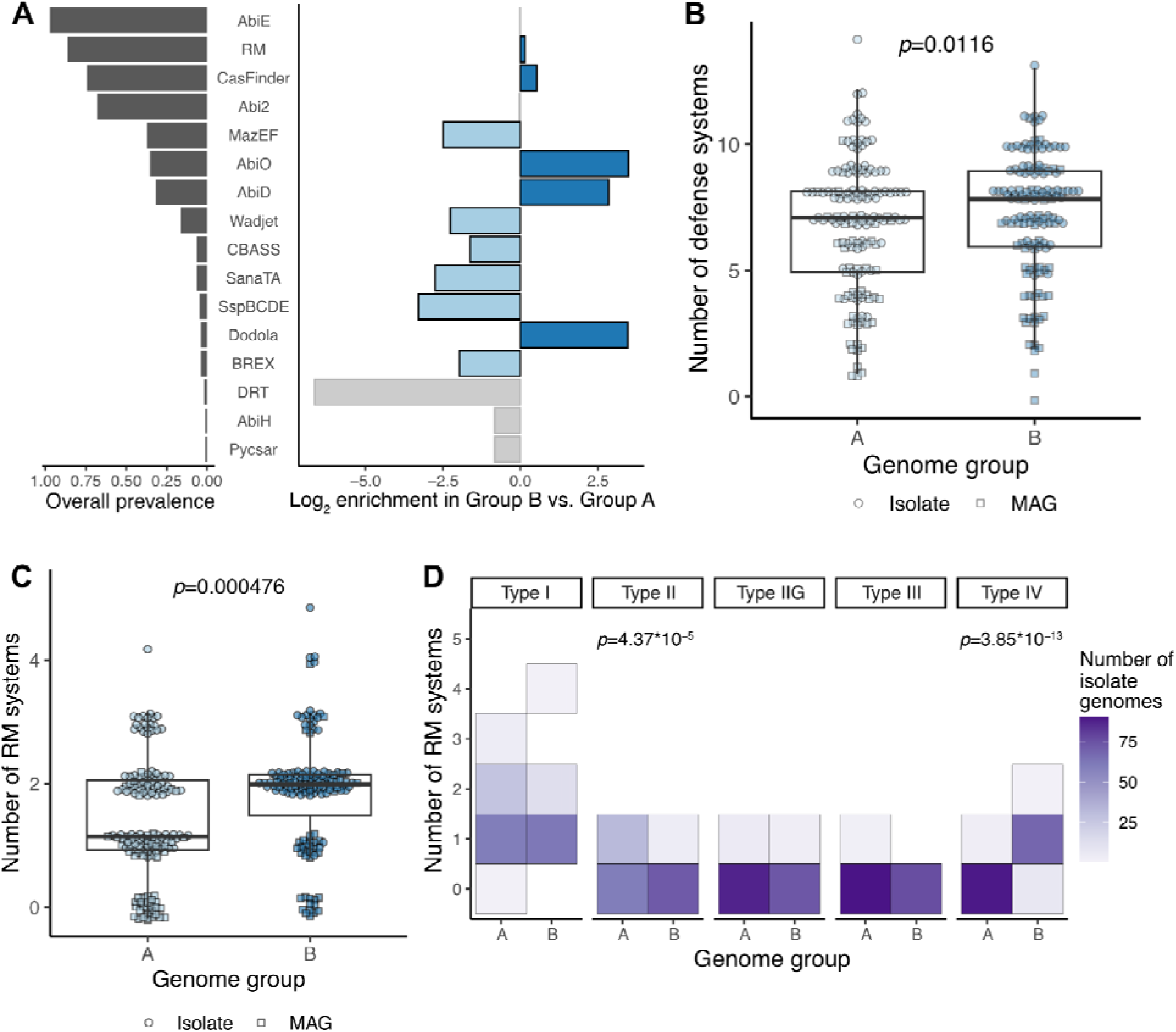
*E. lenta* defense systems are varied and distinct between lineages. **(A)** Left graph, prevalence of defense system types in the *E. lenta* pangenome (isolates and MAGs; *n=*284, DefenseFinder). Right graph, differential abundance in Group B vs. Group A (blue indicates FDR-adjusted *p*<0.05, logistic regression model accounting for genome type (isolate or MAG) and genome size). **(B,C)** Group B has an elevated number of defense systems **(B)** and restriction-modification systems **(C)**. Points represent individual genomes (*n*=284). **(D)** The types of restriction-modification systems present differ between genome groups (*n=*116 isolate genomes). The heatmap is a 2-dimensional histogram indicating the number of genomes with the corresponding RM system copy number. **(B-D)** *p*-values indicate the group effect in Poisson regression models adjusting for genome type (isolate or MAG) with natural log genome size as an offset.

Particularly in light of this divergence in defense systems, we wondered whether different lineages of *E. lenta* might also harbor different sets of plasmids. We identified putative plasmids in our complete hybrid genomes based on circularity, size, and sequence markers (**Methods**). We found a total of 169 mobile elements, including 3 clusters of closely related plasmids detected in at least 10 genomes (**Figures 5A, S5A-F, and Table S5**). As expected, plasmids were detected more frequently in isolate genomes than in MAGs, but their size and gene content were similar across both genome types, supporting our approach (**Figure S5A-D**).

**Figure 5.**
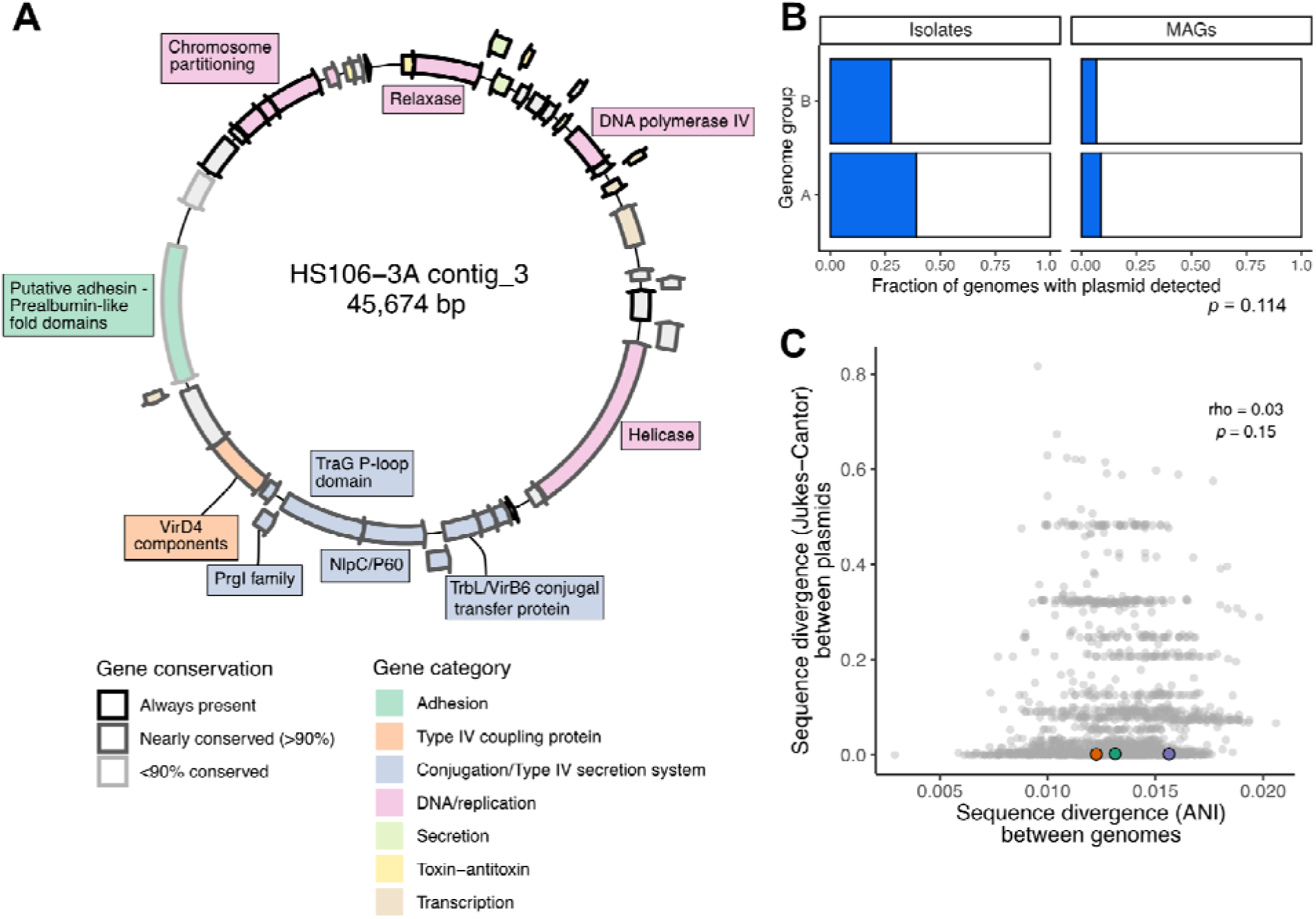
Discovery of a conjugative plasmid common across the *E. lenta* species. **(A)** Gene map of a representative plasmid sequence from *E. lenta* isolate HS106, colored by gene category and shaded by conservation across strains (Bakta/eggNOG annotations). **(B)** The conjugative plasmid is detectable in both *E. lenta* genome groups (*p*-value, genome group in a logistic regression model adjusting for genome type (isolate or MAG; *n=*284 genomes). **(C)** Bacterial host and plasmid sequence divergence are not significantly correlated (*p-*value, Spearman rank-based correlation). Dots indicate pairwise comparisons between plasmid-containing genomes (*n=*66), with colors indicating multiple strains isolated from the same individual.

The most common mobile element was equally common in Group B as in the rest of the species (**Figure 5A,B**). It was large in size (44-48kb) and found in 23.2% of genomes in our collection, across all continents and data sources in this study. It included the major components of Type IV conjugative plasmids in Gram-positive bacteria: genes for mobilization (relaxase), rolling circle replication, and structural and assembly components of the secretion system including multiple transfer protein homologs (**Figure 5B, Table S5**). However, it did not encode a complete set of traditional conjugative pilus structural genes, as has also been observed for similar elements in other Gram-positive species^43^.

This plasmid did not appear to carry major cargo genes in the traditional categories of antimicrobial resistance or known virulence factors, but it did include two potential cargo genes. One of these was a large (2,211 amino acid) helicase gene with a possible role in phage defense or nucleic acid maintenance, as it also has similarity with adenine methylase enzymes. Another candidate cargo gene was a large putative adhesin with similarity to the epithelial-binding SpaA-encoded pilin of *Corynebacterium diphtheriae* (found on 63/66 plasmid contigs), which could act as a putative colonization factor. This gene varies in structure between instances of the plasmid, but its most commonly appearing form is 1,411 amino acids long and contains a transmembrane domain on the C-terminus and 6 repeated SpaA domains. Interestingly, *E. lenta* genomes in our dataset carried as many as 9 genes in this adhesin superfamily (COG4932, median of 3). Adhesin genes vary widely in their domain structure and appear to have multiple evolutionary origins (**Figure S5G**).

We found evidence of potential plasmid transmission in our isolation cohort. Among subjects from whom we obtained 2 or more distinct *E. lenta* isolates (*n*=5 with the plasmid), 3 had multiple strains carrying the plasmid with near-100% sequence similarity (**Figure 5C**). In two of these individuals, the plasmid was shared between a Group B isolate and a Group A isolate. These observations are supportive of active or recent exchange of this plasmid between distinct strain populations, including across lineages.

Other common plasmids were generally small, lacking obvious selective markers, and distributed across the species (**Figure S5C,D**). These plasmids most commonly carried a replication protein, one or more transcriptional regulators, a potential toxin-antitoxin system, and one or two genes of unknown function (**Table S5**). Notably, none of the mobile elements we identified encoded any anaerobic respiratory reductases, suggesting that these strain-variable genes may be evolving by mechanisms other than plasmid-mediated gene transfer.

### *E. lenta* Group B has distinct associations with inflammatory bowel disease

Finally, we hypothesized that our expanded dataset of high-quality reference genomes could provide a basis for improved *E. lenta* strain tracking in metagenomic samples. Since Group B comprises a single clade, we focused on quantification of this group, testing both single nucleotide polymorphism (SNP)-based and gene-based methods. Briefly, we defined a set of common SNPs in *E. lenta* core genes that distinguish the Group B lineage from other *E. lenta* (**Figure S6A,B, Methods**), and we also compared the presence and abundance of Group B-linked marker genes (**Figure 3, Table S3**)^44^. We applied these methods first to metagenomic data from the ImmunoMicrobiome cohort^31^, validating that the overall *E. lenta* abundances estimated by our core gene set were consistent with our isolation results (**Figure S6C, Methods**) and highly correlated with published methods (**Figure S6D**). Group B SNP-based abundances were also highly correlated with the abundances of Group B-linked gene families (**Figure S6E**). Consistent with our isolation results, total *E. lenta* core gene coverage was often greater than Group B coverage, indicating potential coexistence of multiple *E. lenta* lineages (**Figure S6F**).

Given *E. lenta*’s documented effects on host immune signaling^23,25^, we wondered whether the Group B lineage differs from the rest of the species in its associations with immune cell populations. We took advantage of immune profiling data (CyTOF) generated from healthy subjects in the ImmunoMicrobiome cohort study^31^ from blood samples provided at the same time as the stool samples used for isolation. However, despite the high prevalence of *E. lenta* in this cohort, neither total *E. lenta* nor Group B abundances were significantly associated with any immune population after false discovery rate correction (**Figure S7A-B**).

We then quantified *E. lenta* lineage abundances in a disease context, using published metagenomic datasets from patients with inflammatory bowel disease (IBD) and healthy controls and applying strict quality cutoffs (**Methods**)^7,45,46^. As expected^23^, the total abundance of *E. lenta* was elevated in IBD patients compared with controls (**Figure 6A**). Group B was similarly increased in patients compared with controls (**Figure 6B**). Further, Group B-linked gene family abundances, including the previously identified nitroreductase operon (**Figure 3B**), were more strongly associated with ulcerative colitis than *E. lenta* core genes (**Figure 6C**). This pattern was also observed albeit to a lesser degree in Crohn’s disease patients (**Figure 6D**).

**Figure 6.**
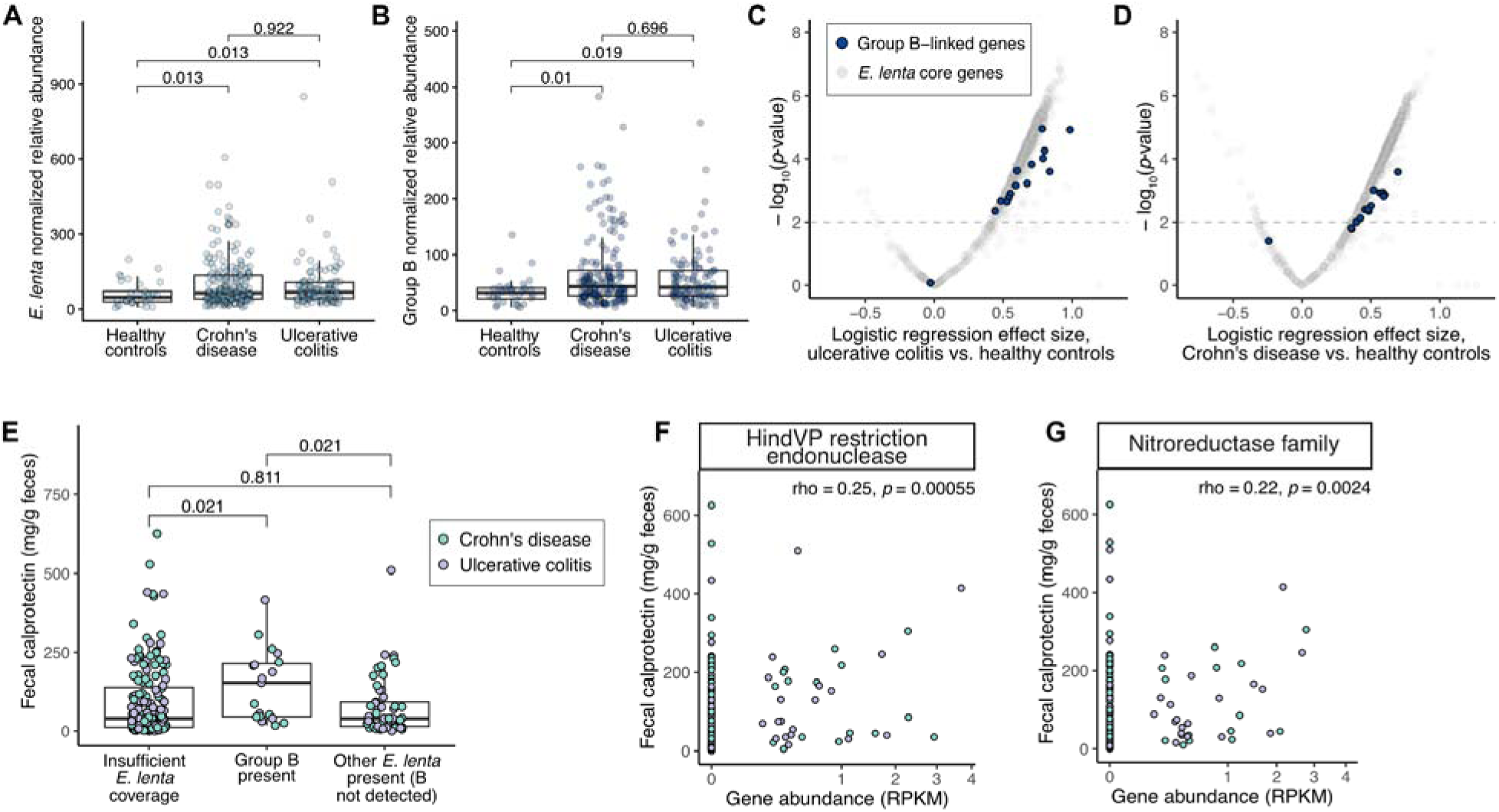
*E. lenta* Group B is associated with inflammatory bowel disease severity. **(A,B)** Both total *E. lenta* **(A)** and *E. lenta* Group B **(B)** are significantly enriched in samples from subjects with IBD (UC, ulcerative colitis; CD, Crohn’s Disease) relative to healthy controls (HC). Abundance is shown in units of average sample coverage of core gene SNP sites per billion reads (see **Methods**). **(C,D)** Group B-associated gene abundances (RPKM) are enriched in subjects with UC and CD. Points indicate gene families (*p-*value, logistic regression model of disease status with study as a covariate, *n=*231 HC, *n=*320 UC, *n=*601 CD). Gray indicates the core genome (>95% of genomes) and blue indicates the 16 gene families most strongly linked with Group B (Figures 3**, S4, and Table S4**). (**E)** Fecal calprotectin is higher in samples positive for *E. lenta* Group B. Green (*n=*144 CD with calprotectin data), grey (*n=*89 UC). One outlier sample with insufficient coverage and a calprotectin measurement of 2,440 mg/g feces is not shown but was included in statistical testing. **(A,B,E)** *p*-values, Dunn’s test with multiple hypothesis correction. **(F,G)** Normalized abundances (reads per kilobase per million reads, RPKM) of two representative gene families enriched in Group B that are significantly correlated with fecal calprotectin (green *n=*144 CD, grey *n=*89 UC). Two outlier samples with calprotectin > 2,000 mg/g feces are not shown but were included in statistical testing. Spearman correlation coefficients and *p-*values are shown. Gene abundances are on a square root scale for better visibility of low abundances.

Notably, *E. lenta* Group B was also associated with fecal calprotectin, a marker of gut inflammation. Both the presence of Group B (**Figure 6E**) and the abundance of Group B-linked genes (**Figure 6F,G**) were linked with greater levels of calprotectin. The association with Group B presence persisted after accounting for diagnosis and study as covariates (*p*-value=0.05). Overall, these results suggest that *E. lenta* Group B is widespread in IBD patients and particularly elevated in the context of active inflammation.

## DISCUSSION

In this study, we used a novel selective media formulation to greatly expand the available set of *Eggerthella lenta* cultured isolates and high-quality genomes. One previous study has reported selective isolation of *E. lenta* using the antibiotics cefotaxime and ceftriaxone as selective agents^47^; however, additional microscopy and PCR screening was required to identify *E. lenta* colonies. We used a simple chemically defined media, ESM, which enables sensitive, specific, and quantitative isolation of *E. lenta* from human stool samples. Compared to culturomics approaches that have increased the number of available *E. lenta* isolates using microfluidics and/or robotics^48,49^, plating on ESM agar is technically straightforward and low-cost. The new isolate genomes obtained from our cohort have nearly doubled the number of publicly available high quality *E. lenta* isolate genomes.

Using our new media formulation, we detected *E. lenta* in a cohort of North American adults with 90.1% prevalence (82 out of 91 samples). This prevalence highlights the broad distribution of this bacterial species and is consistent with previous qPCR-based findings that *E. lenta* is often undersampled by metagenomics^29^. Across our broader collection, all but 4 genomes of known origin (98.5%) came from North America, Europe, or China. This bias could indicate high prevalence of *E. lenta* in these regions compared to others, but it is also likely influenced by their overrepresentation in deep microbiome sequencing datasets^50,51^. Further sampling and more quantitative analyses will be needed to resolve these effects.

The *E. lenta* pangenome is large, variable, and minimally associated with geography. This low level of phylogeographic signal is concordant with a recent large study of phylogeography in gut microbial species, which placed *Eggerthella* in the bottom third of taxa by strength of phylogeographic signal^52^. Furthermore, we observed frequent coexistence of multiple *E. lenta* lineages within the same host in our cohort (*n=*6 subjects, **Table S2**). Together, these findings could indicate sympatric diversification of *E. lenta* into distinct niches structured by phage defense, use of electron acceptors, and stress tolerance, rather than by geographic isolation. In particular, the extensive strain-level differences in phage defense genes emphasize the importance of phages as a driver of strain diversity, a pattern that has been observed in other gut bacterial species^53,54^.

We identified and characterized a dominant lineage with a reduced genome within the *E. lenta* species, termed Group B. This group displayed clear functional differences from the rest of the *E. lenta* species, including distinct sets of respiratory enzymes, an enrichment in phage defense systems, and potential differences in stress response. The phage defense system linked to Group B included a putative Type II restriction-modification system, a Type IV restriction enzyme, and a large helicase domain-containing gene. The combination of a cytosine methyltransferase and a large helicase/nuclease is reminiscent of the recently-described Druantia anti-bacteriophage defense system^55^, although the domain structure and operon organization do not match any described subtype.

Group B’s tolerance to nitric oxide and positive association with fecal calprotectin could suggest specialization of this lineage to a niche hostile to many other gut microorganisms. Elevation of nitric oxide and other free radical compounds is a known feature of the inflamed gut^56^, with the capability to shape microbiota composition^57^. Notably, Group B most likely represents a different lineage than the clade described by Kumbhari *et al*.^7^, which was reported to have a negative association with calprotectin based on much of the same metagenomic data. These opposite lineage-specific associations are consistent with the known ability of *E. lenta* strains to catalyze metabolic reactions that may mitigate^25^ or exacerbate^23^ inflammatory bowel disease. As the number of *E. lenta* isolates and high-quality genomes continues to grow, we anticipate that future studies will continue to define *E. lenta* subpopulations and their potential positive and/or negative impacts on host immunity.

We also uncovered a putative conjugative plasmid, broadly shared across multiple *E. lenta* lineages. This discovery is promising for the advancement of genetic tool development in the *Eggerthellaceae,* which remains technically challenging^58^. Conjugation-based approaches have enabled effective genetic modification in several other Gram-positive gut bacterial taxa^59–62^. The prevalence of this plasmid indicates the potential for similar success in *E. lenta*.

Another notable aspect of this conjugative plasmid was the presence of a large adhesin gene. A gene in the same superfamily (COG4932, Clumping factor A-related surface protein) was previously negatively associated with colonization success in a competition experiment in which 22 strains of *E. lenta* were co-inoculated into gnotobiotic mice^29^ (**Figure S5G**). It is possible that this conjugative plasmid may contribute to variability in surface interactions of *E. lenta* and that these adhesin genes could influence fitness in heterogeneous environments. Genes encoding colonization factors have been commonly found on conjugative plasmids in rhizobia, impacting the outcome of their symbioses with legumes^63^.

To date, *E. lenta* strain variation has typically been described in terms of a small number of genes involved in specialized metabolism and based on a relatively small number of genomes^29^. Our collection provides a broader context for this work while highlighting how much remains unknown. For example, we found that some of *E. lenta*’s specialized enzymes could be linked to a specific lineage, whereas others were broadly distributed, enabling follow-on studies aimed at identifying the mechanisms underlying variation in key gene families relevant to host pathophysiology. Importantly, our selective media and analysis framework will enable continued sampling of *E. lenta* across broader cohorts, helping to further clarify the factors shaping *E. lenta*’s multifaceted roles in the human gut across populations.

## RESOURCE AVAILABILITY

### Lead contact

Further information and requests for resources and reagents should be directed to the Lead Contact Peter Turnbaugh.

### Materials availability

Bacterial strains isolated as part of this study will be made available from the authors upon reasonable request.

### Data and code availability

Genome data has been deposited in NCBI BioProject Accession Pending. Metagenomic data is available from NCBI BioProject PRJNA1390888 (ImmunoMicrobiome study) and from inflammatory bowel disease cohorts via NCBI BioProject IDs PRJNA993675 and PRJNA398089 and the Human Microbiome Bioactives Resource Data Portal. Scripts used to generate all figures are available at www.github.com/cnoecker/ElentaGenomes. Genomes, gene annotations, and associated metadata for the de-replicated set of 284 *E. lenta* genomes are also available from Zenodo (doi:10.5281/zenodo.19325436).

## ACKNOWLEDGMENTS

We acknowledge Jordan Bisanz, Margaret Alexander, and other current and former Turnbaugh lab members for helpful suggestions and feedback, Katherine Xue for advice on metagenomic strain profiling, Daryll Gempis for assistance with selective media preparation, and Brittany Davidson for assistance with ImmunoMicrobiome study samples. We thank all the study participants of the ImmunoMicrobiome cohort. We thank Kenya Honda and Koji Atarashi for sharing genome assemblies, and Harris Wang and Yiming Huang for sharing *Eggerthellaceae* isolates.

We acknowledge the Benioff Center for Microbiome Medicine for assistance with nanopore sequencing. We acknowledge the Minnesota Supercomputing Institute (MSI) at the University of Minnesota and the Wynton HPC Co-Op cluster at UCSF for computing resources. Funding was provided by the National Institutes of Health (R01CA255116, R01DK114034, and R01HL122593 to P.J.T.; F32GM140808 to C.N.; R01DE032033 and R01CA290027 to M.H.S.), the UCSF Benioff Center for Microbiome Medicine (Trainee Pilot Award to C.N. and J.B., research funding to M.H.S.), the National Science Foundation (award #2437513 to C.N.), DOD US Army Med. Res. Acq. Activity (award BC220499 to M.H.S.), the Bakar ImmunoX Initiative (to M.H.S.), and Minnesota State University, Mankato (Faculty Research Grant to C.N. and Undergraduate Research Center grants to C.D. and N.R.). P.J.T is a Biohub, San Francisco, Investigator. Illumina sequencing was performed at Biohub, San Francisco and the UCSF Center for Advanced Technology.

## AUTHOR CONTRIBUTIONS

Conceptualization: C.N. and P.J.T.; Data curation: C.N.; Formal analysis: C.N., L.G., C.D., N.R., and F.D.; Funding acquisition: C.N., J.B., M.H.S., P.J.T.; Investigation: C.N., L.G., C.D., N.R., F.D., L.R.H, T.S.K., K.T., J.B.; Methodology: C.N., L.R.H., C.W.Y.H., C.L.G., P.J.T.; Project administration: C.N.; Resources: C.N., J.B., M.H.S.; Software: C.N.; Supervision: C.N., M.H.S., P.J.T.; Validation: C.N.; Visualization: C.N., L.G., C.D., N.R., F.D.; Writing - original draft: C.N. and P.J.T.; Writing - review and editing: C.N. and P.J.T.

## METHODS

### Isolation of *E. lenta* from human stool samples

Stool samples from the ImmunoMicrobiome cohort study^31^ (collected under UCSF IRB approved protocol #18-27022) were aliquoted into pre-weighed tubes on dry ice in a biosafety cabinet and stored at -80°C. A stool slurry was prepared aerobically from a single aliquot of each sample as follows. We prepared a sterile solution of BHI^CHV^: BHI broth (BD Bacto #237500) supplemented with 0.05 g/L cysteine, 1 mg/L hemin, and 5 mg/L menadione (vitamin K_3_ suspended in ethanol). 15% sterile glycerol was added to the media and the sample was then diluted at 1 g sample per 10 mL media. The tube was thoroughly vortexed until the sample was fully suspended, after which particles were allowed to settle for 5 min and 500 μL of supernatant transferred to a fresh tube. Stool slurries were transferred to a Coy Laboratory Products Type B anaerobic chamber filled with <5% hydrogen, 20% carbon dioxide, and nitrogen mixed gas.

Our recipe for *Eggerthella* selective media (ESM) is found in **Table S1**. Typically, a 2X solution of most components was prepared and sterilized with a 0.22 μm vacuum filter, after which it was supplemented with sterile solutions of vitamins and trace minerals. Trace solutions were prepared and stored as described previously^30^. A 2x solution (30 g/L) of noble agar or bacteriological agar was prepared, autoclaved, and allowed to cool to approximately 55°C. The agar solution was then combined with the filter-sterilized 2X media, stirred, and used to pour plates.

The selectivity of ESM was validated on 3 randomly selected stool samples. Serial dilutions of 10^−4^, 10^−5^, and 10^−6^ were prepared, and 1 mL of diluted inoculum plated onto ESM agar using large 150 mm Petri dishes prior to anaerobic incubation at 37°C for 7 days (*n*=3 replicates/dilution). We counted colonies from all plates for estimation of colony-forming units. A selection of colonies from at least one replicate plate were selected for streak purification. All remaining plate biomass was scraped into 400 μL sterile deionized water using a sterile plastic rod and stored at -80°C, pooling the material from all 3 replicate plates into a single tube for each sample and dilution.

Based on this pilot experiment, we opted to dilute all remaining slurries 10^−5^ in sterile pre-reduced PBS using serial dilution. In two cases, 10^−3^ and 10^−4^ dilutions were plated after no colonies were obtained at 10^−5^, yielding a single isolate from each (indicated in **Table S2**). We plated 400 μL of each diluted inoculum on ESM agar plates and then spread the sample using sterile glass beads (*n*=3 plates/sample). We then dried each plate for 5-10 min, removed the beads, and incubated them at 37°C for 7 days.

We selected at least 1 colony per plate for two rounds of streak-purification on rich media (BHI^A^, BHI agar with 1% arginine). For plates with multiple distinct or ambiguous colony morphologies, one colony of each morphology was carried forward. 1-2 additional rounds of streak purification were performed as needed to remove apparent contamination. Taxonomic identity was confirmed using full-length Sanger 16S rRNA gene sequencing using the standard primers 8F (5’-AGAGTTTGATCCTGGCTCAG-3’) and 1542R (5’-AAGGAGGTGATCCAGCC-GCA-3’). All but one colony were confirmed as *E. lenta*. We also obtained and cultured 5 *Eggerthellaceae* isolates obtained from a recent culturomics study^48^. These were received as glycerol stocks and processed for DNA extraction and long- and short-read sequencing as described below.

### Testing of bacterial isolate growth in ESM

Bacterial glycerol stocks were streaked onto BHI agar supplemented with 1% arginine, 0.05% L-cysteine-HCl, 1 μg/mL vitamin K, and 5 μg/mL hemin, and incubated anaerobically for 3 days at 37°C. At that time, a single colony from each strain was inoculated into 5mL of BHI+ broth. After 48 hours of growth, strains were washed anaerobically in phosphate-buffered saline and resuspended to a final OD of ∼0.1. 20 μL of the resuspended cells were used to inoculate a 96-well plate (Corning) containing 180 μL of reduced ESM broth. The plate was incubated at 37°C in a Biotek Eon plate reader for 72 hours. Every 30 minutes, 40 seconds of shaking was performed prior to measuring absorbance at 600 nm.

### DNA extraction and sequencing

The pooled biomass collected in the pilot experiment was used for amplicon sequencing of the V4 region of the 16S ribosomal RNA gene using a two-stage amplification^64^. Briefly, DNA was extracted using the ZymoBIOMICs 96 MagBead DNA Kit, following the manufacturer protocol with homogenization by 5 minutes bead beating (BioSpec Mini-Beadbeater-96), followed by 5 min room temperature incubation, and a repeat of 5 min bead beating. Samples were amplified with primers 515F (5’-GTGYCAGCMGCCGCGGTAA-3’) and 806R (5’GGACTACNVGGGTWTCTAAT-3’) using a reaction mix of 0.45 μL DMSO, 0.0045 μL SYBR Green I 10x diluted in DMSO to 1000x, KAPA HiFi PCR kit (#KK2502, 1.8 μL 5x KAPA HiFi Buffer, 0.27 μL 10 mM dNTPs, 0.18 μL KAPA HiFi polymerase), 0.045 μL of each amplification primer (final concentration 1 μM), 6.2055 μL nuclease-free water, and 1 μL DNA. A BioRad CFX 384 real-time PCR instrument amplified four 10-fold serial dilutions of DNA with the following parameters: 5 min 95°C, 20x (20 sec 98°C, 15 sec 55°C, 60 sec 72°C). Non-plateaued individual sample dilutions were selected for secondary indexing PCR. In the secondary reaction, the KAPA HiFi PCR kit was used with: 4 μL 5x KAPA HiFi Buffer, 0.6 μL 10 mM dNTPs, 1 μL DMSO, 0.4 μL KAPA HiFi polymerase, 4 μL indexing primer, 10 μL of 100-fold diluted primary PCR reaction, with identical amplification parameters as above. Amplicons were quantified with PicoGreen according to manufacturer’s instructions, equimolar pooled, and gel purified (QIAquick Gel Extraction Kit). Libraries were quantified with KAPA Library Quantification Kit for Illumina Platforms according to the manufacturer’s instructions, spiked with 15% PhiX, and sequenced on an Illumina MiSeqV3 instrument. Primers and adapters were removed using the *cutadapt trim-paired* command in QIIME2^65^ (v2020.11). Sequences underwent trimming to 220 bp (forward) or 150 bp (reverse), quality filtering, denoising, and chimera filtering using DADA2^66^ (v1.18.0) with QIIME2 command *denoise-paired*. Sequence length was filtered to 250–255 bp with QIIME2 command *feature-table filter-seqs*. Taxonomy was assigned to amplicon sequence variants (ASVs) using the SILVA v138 database^67^.

High-molecular weight DNA was extracted from *E. lenta* isolates using a modified version of the Qiagen MagAttract High-Molecular Weight DNA Kit (#67563) to maximize yield from difficult to lyse bacterial cells. A single colony of each *E. lenta* isolate was grown in 5 mL BHI^A^ for 40-48 hours (OD_600_ 0.7-1.2). We transferred 1.8 mL of each culture to a 2mL Eppendorf tube and centrifuged it 4 min at 10,000 x *g*. The supernatant was removed with a pipette and pellets were either frozen at -80°C until extraction or extracted immediately. We resuspended each cell pellet in 450 μL Buffer P1 and then transferred the resulting liquid into a FastPrep Lysing Matrix E tube (#1169140-CF) prior to homogenization in a Biospec Mini-Beadbeater-96 (#1001) for 30 seconds. Tubes were centrifuged very briefly (15-30 seconds at 7,000 rpm), after which 250 μL of each supernatant was transferred to a fresh low-adhesion microcentrifuge tube (USA Scientific #1415-2690). 50 μL of 100 mg/mL lysozyme (Roche, #10837059001) was added to each tube. Lysates were incubated at 37°C for 2 hours on an Eppendorf Thermomixer shaking at 900 rpm. After incubation, we used the manufacturer’s protocol for magnetic bead-based DNA purification. DNA was eluted into 100 μL of Qiagen Buffer EB, transferred into a fresh low-adhesion tube using wide bore pipette tips to prevent shearing, and stored at 4°C. We quantified DNA concentration using a Qubit High Sensitivity DNA assay (#Q33231) and size distribution using an Agilent Tapestation with a Genomic DNA ScreenTape (#5067-5365) using the manufacturer’s protocols.

We prepared genomic DNA libraries for long-read sequencing using the Oxford Nanopore V14 (SQK-NBD114-24) or V12 (SQK-NBD112-24) kits (see **Table S2**), using the manufacturer’s protocol as provided in May 2023. Sequencing was performed on the corresponding flow cell (R10.4.1 V14, R10.4 V12) on a MinIon Mk1B instrument, following the manufacturer’s protocol. We loaded 10-24 genomes per flow cell, which then underwent a 72-hour sequencing run. The same extracted DNA was used for library preparation for Illumina sequencing. A library was prepared using the Illumina DNA Prep Kit, following a modified version of the manufacturer’s protocol with reduced volumes per sample (as described at dx.doi.org/10.17504/protocols.io.be6rjhd6). Samples were amplified using either the EPM master mix included in the kit or the Kapa HiFi HotStart ReadyMix (#KK2612) with custom 12 base pair dual indexes provided by Biohub, San Francisco. Samples were sequenced on an Illumina NextSeq 2000 P3 at Biohub, San Francisco.

### Sequence data processing and genome assembly

Nanopore data (POD5 files) were basecalled and demultiplexed using guppy (v6.5.7) models on a GPU with the “high-accuracy” setting and chunk sizes of 500 reads. Reads were assembled with Flye^68^ (v2.9.2) with default parameters with the addition of the “--plasmids” flag and an estimated genome size of 3.6Mb. Assemblies were then polished using Medaka (v1.8.0, https://github.com/nanoporetech/medaka) with the consensus model chosen based on the nanopore flow cell used (10.4 or 10.4.1). Polished assemblies were coordinate-adjusted using the *fixstart* command in Circlator^69^ (v1.5.5). Short (Illumina) reads were then aligned to the resulting genome using *bwa mem*^70^ (v0.7.17), and the alignment was used for a second polishing with Polypolish^71^ (v0.5.0). In cases where the longest nanopore-only contig obtained by Flye was less than 1 Mb (typically indicating that limited long-read data was available), Unicycler^72^ v0.5.0 was instead used to perform an Illumina-first assembly with scaffolding assistance from the nanopore data. For these genomes (indicated in **Table S2**), polishing and coordinate adjustment was not performed.

### Compilation of *Eggerthellaceae* genomes and metadata

We obtained a total of 116 *E. lenta* genomes, including 111 from ImmunoMicrobiome sample isolations and 5 previously published isolates^48^ (**Table S2**). We supplemented these with genomes spanning the broader *Eggerthellaceae* family compiled across multiple public repositories and research groups (**Table S3**). This includes 1,445 genomes labeled as *Eggerthellaceae* in NCBI Genome (July 2024), excluding those marked as atypical, and 1,959 genomes labeled as *Eggerthellaceae* from the United Human Gastrointestinal Genome (UHGG) catalog (v2.0.1 from MGnify^73^). We also included 6 genomes from a recent study that were not deposited in NCBI or UHGG^48,74^. The *Eggerthella lenta* ATCC 43055 assembly was downloaded from the ATCC Genome Portal (genomes.atcc.org). Metadata for NCBI genomes was obtained from the BioSample database (using the *biosample2table* function from https://github.com/stajichlab/biosample_metadata). We then merged and curated this metadata with the provided metadata for UHGG^75^, the ImmunoMicrobiome cohort^31^, and the indicated papers^48,74^. The *countrycode* R package was used to assign continents.

We performed multiple quality control steps to ensure that only high-quality complete genomes were included in the final analysis. GTDB-tk v2.3.2^76^ was first used to verify that all of the genomes are from the *Eggerthellaceae* family, resulting in the removal of 79 misclassified genomes, all from NCBI. We then used GUNC v1.0.6^77^ (default parameters, proGenomes database) to identify genomes with a passing score and a combination of CheckM2 v1.0.2^78^ and BUSCO v5.7.1^79^ (using Coriobacteriia marker genes) to identify genomes with a consistent completeness >80% and contamination <5%. We used a completeness threshold of 80% rather than the typical threshold of 90% for high-quality MAGs in order to maintain geographic diversity in our dataset (since most genomes not from North American hosts were MAGs). Finally, the 1,991 genomes that passed all of these quality control steps were dereplicated at 99.9% nucleotide identity and 95% aligned fraction using skDER v1.2.2^80^, which selects genome representatives from closely related clusters based on contiguity (N50) and connectedness (similarity to the largest number of genomes). In total, these analyses resulted in a final set of 1,425 high-quality dereplicated *Eggerthellaceae* genomes, including 284 assigned to the *E. lenta* species (**Table S3**).

### Gene annotation

We annotated the 1,425 high-quality dereplicated genomes (**Table S3**) with Bakta v1.9.3^81^ (default bacterial parameters) to identify open reading frames and RNA genes and with DefenseFinder v1.2.2^38^ with default parameters to identify antiviral defense systems. We then annotated open reading frames with eggnog-mapper v2.1.2^32^ (setting the *tax_scope* flag to “bacteria”) to identify eggNOG orthologous gene families and secondly with Interproscan v5.69.101.0^82^ to identify protein domains with default parameters. To define reductase gene families at a higher resolution than their eggNOG classification, all amino acid sequences assigned by eggnog-mapper to the COG superfamilies of COG1053 (flavin-dependent reductases) and COG0243 (molybdopterin-dependent reductases) were sub-clustered using ProteinOrtho v6.3.4^83^ with 90% sequence similarity and 80% coverage. We also used the *fast.genomics* web tool^84^ (accessed December 2025) to search for homologs of the gene families in the nitroreductase operon in diverse bacteria. Gene regions were visualized using the *gggenes* v0.5.1 and *gggenomes* v1.0.1^85^ R packages.

### Plasmid identification, annotation, and clustering

We identified plasmid contigs initially only in our newly sequenced hybrid assemblies, using a combination of methods. First, we profiled contigs with GeNomad v1.8.0^86^, which identifies likely plasmids and viral elements. We next ran CircleMap v1.1.4 on short-read alignments to our hybrid genomes, which identifies circular contigs with an associated confidence score. We defined likely plasmid contigs as those that had a plasmid score (GeNomad) >0.5, a circularity score (CircleMap) >100, and a sequence length <200 kbp, based on high agreement between these tools at these cutoffs and examination of the distributions of scores across all contigs. We assigned genes on these contigs to high-resolution gene families using ProteinOrtho (90% identity, 80% coverage), and grouped similar sequences into clusters of similar sequences using hierarchical clustering based on gene family presence/absence (average linkage, cut at a height of 0.5). Lastly, we used BLASTn v2.13.0 to search for long sequences similar to these in the broader dataset, filtering for those with 100% query coverage and at least 50% nucleotide identity with a likely plasmid.

We performed additional annotation analyses of plasmid contigs to uniformly group and compare these sequences. We reoriented conjugative plasmid sequences to begin at the site of a DNA polymerase gene, if present, in order to have consistent coordinates across closely related plasmids. Plasmids were then re-annotated with Bakta to obtain annotations with these consistent coordinates and detect gene fragments that might have been fragmented at the beginning/end of the sequence. To supplement this annotation, we also used the web interface of oriTfinder^87^ to confirm annotation of conjugation genes. Plasmid maps were generated using a modified version of functions from the *plasmapR* package (https://github.com/BradyAJohnston/plasmapR).

### *E. lenta* comparative genomics

Pangenome curves were estimated based on presence/absence of eggNOG orthologous gene families (annotated by eggnog-mapper) in repeated random sampling of genome subsets. To assess the openness or closeness of the pangenome, a Heaps’ law curve was fit^88^ using the *heaps* function implemented in the R package *micropan* v2.1 with 500 permutations^89^. As a supporting pangenome analysis, we used the Panstripe package v0.3.3^90^ with the default parameters to fit a model of gene gain and loss, accounting for phylogenetic relatedness between genomes.

To assign the 284 *E. lenta* genomes to clusters (genome groups), we used a Mash distance approach^91^, as described^13^. We calculated pairwise Mash distances and then hierarchically clustered the resulting distance matrix using Ward’s method. To evaluate the appropriate number of clusters (i.e. dendrogram cut height), we generated elbow and silhouette plots as implemented in the *factoextra* R package v1.0.7^92^, leading us to cut the cluster dendrogram at a height of 0.06. We also assigned genomes that were not included in the final dereplicated set to Group A or Group B based on the assignment of their three nearest neighbors in terms of Mash distance (**Table S3**). If there was not a unanimous assignment across the 3 neighbors, the genome was left as ambiguous.

Whole-genome average nucleotide identity was also calculated using PyANI v0.2.13.1^93^. We constructed a phylogeny of all *E. lenta* genomes using Phylophlan v3.1.1^94^ with the *E. lenta*-specific marker database, amino acid sequence alignment, and the low-diversity setting*. Eggerthella timonensis* Marseille-P3135 was included as an outgroup. A supporting phylogenetic tree was also constructed based on a core gene alignment inferred by Panaroo v.1.5.0^95^ using RAxML v8.2.12^96^.

We identified gene families associated with *E. lenta* Group B using two complementary methods. For both methods, we included eggNOG gene families present in greater than 10% and less than 90% of genomes. First, we implemented a logistic regression model that accounted for Group B membership, genome size, and genome type (either isolate or MAG) (i.e. *GenePresence ∼ GroupBStatus + GenomeSize + GenomeType*). Second, we used a machine learning-based classification approach, building a random forest model to predict Group B membership based on eggNOG gene family presence/absence. The model was fit with the R package *ranger* v0.17.0^97^ with the default parameters. Permutation-based importance scores were calculated with *ranger* for each feature included in the model. The set of 16 most strongly linked genes were identified as those with a logistic regression coefficient greater than 4.5 and/or a random forest permutation-based importance greater than 0.004, based on agreement between both methods at these cutoffs and the observation that most of the genes that met either threshold were co-located in Group B genomes (**Figure S4**).

### Nitrosative stress assay

A 5 mM stock solution of diethylamine NONOate (DETA NONOate, MedChemExpress #HY-136278) was prepared in ultrapure water, filter-sterilized with a 0.22 μM syringe filter, and transferred into 0.5 mL aliquots on ice. Aliquots were stored at -80°C and a fresh aliquot was used for each experiment. *E. lenta* strains were streaked on BHI^A^ agar and incubated anaerobically for 3 days at 37°C. At that time, a single colony from each strain was inoculated into 5 mL BHI^A^. To set up the experiment, BHI^A^ was combined aseptically with the DETA NONOate stock to obtain the indicated concentrations at a volume of 1.5 mL for each condition, and then transferred into a 96-well clear round-bottom plate (Corning) with 180 μL per well. The plate was closed and transferred immediately into a Coy Type C chamber with 5% nitrogen, 20% carbon dioxide, and the balance nitrogen. After 24 hours of growth by the starter culture, wells were inoculated with 5 μL of either well-mixed late exponential phase culture or media control (*n*=3-6 replicates/condition). Plates were sealed with a transparent Breathe-Easy sealing gas exchange membrane (RPI) and placed into a Byonoy Absorbance 96 plate reader positioned on a custom adapter on an IKA Twister Stirrer/Shaker (#20027010), all housed in a 37°C incubator in the anaerobic chamber. Absorbance was measured at 600 nm every 20 minutes for 72 hours, with constant shaking at 100 rpm. Growth curves were normalized and analyzed using the *growthcurver*^98^ package v0.3.1. The fraction of maximum growth was calculated for each strain as the median maximum absorbance obtained across replicates at a given NONOate concentration divided by the median maximum absorbance without NONOate.

### Metagenomic sequencing data analysis

We utilized a set of SNPs in core genes to quantify *E. lenta* Group B from multiple metagenomic datasets. First, we used the Panaroo pipeline (v1.5.0)^95^ to define *E. lenta* core genes and construct core gene alignments across all 284 *E. lenta* genomes (**Table S3**). Panaroo-based alignments were filtered to exclude genes with excessive alignment gaps (>30% of positions), revealing 447 core genes. We identified Group B marker SNPs in these core genes in the 95 genomes with near-perfect completeness and contamination, to avoid identification of spurious SNPs (completeness >98% and contamination <2%). Marker SNPs were defined as those found with ≥10% prevalence in Group B but were not seen in Group A. This process resulted in identification of 343 marker SNPs for Group B.

We performed updated taxonomic and functional profiling of the quality-filtered metagenomic data described in the ImmunoMicrobiome study^31^. We performed taxonomic profiling using Metaphlan v4.0.3^99^ and gene family profiling using HUMAnN v3.6.1^44^. Gene family abundances were normalized based on genome equivalents estimated by MicrobeCensus v1.1.1^100^. UniRef90 gene families quantified by HUMAnN were mapped to corresponding eggNOG gene families based on the eggNOG database.

We also analyzed metagenomic data from multiple IBD cohorts, including the NCBI BioProjects PRJNA993675 (PRISM study^46^) and PRJNA398089 (iHMP^101^), and from the Lewis^102^, LSS-PRISM^8^, NL-IBD^46^, RISK^103^, and Stinki^104^ cohorts via the Human Microbiome Bioactives Resource Data Portal (https://portal.microbiome-bioactives.org/), resulting in a total of 1,474 samples with nonzero reads mapped to *E. lenta* core genes, including 287 healthy control and 1,187 IBD patient samples. Associated fecal calprotectin data was obtained from Kumbhari *et al.*^7^ for a subset of samples (*n*=294). Reads were aligned to the *E. lenta* core gene set as described above. Gene family profiling was performed on all samples in this compiled dataset using HUMAnN v3.9 to quantify UniRef90 gene families, which were then mapped to corresponding eggNOG gene families based on the eggNOG database.

Metagenomic sequencing reads were aligned to *E. lenta* core genes using HISAT2 v2.2.1^105^ with the following settings: “*--no-spliced-alignment -p 6 --no-unal --sensitive*”. We analyzed marker sites that had at least 5x coverage of any allele. For *E. lenta* to be called as present with sufficient evidence for lineage analysis, we applied a strict cutoff of detection of at least 21 core genes at the 5x coverage threshold (to limit the influence of off-target alignment). To call Group B present, we required detection of at least 21 core marker genes and at least 8 marker SNPs specific to Group B.

### Quantification and statistical analysis

Unless otherwise specified, all analyses were performed in R v4.4.0 using the packages *data.table*, *ggplot2*, *broom, ggpubr.* Statistical tests were performed in R as described in the figure legends.

## SUPPLEMENTAL FIGURES AND FIGURE LEGENDS

**Figure S1.**
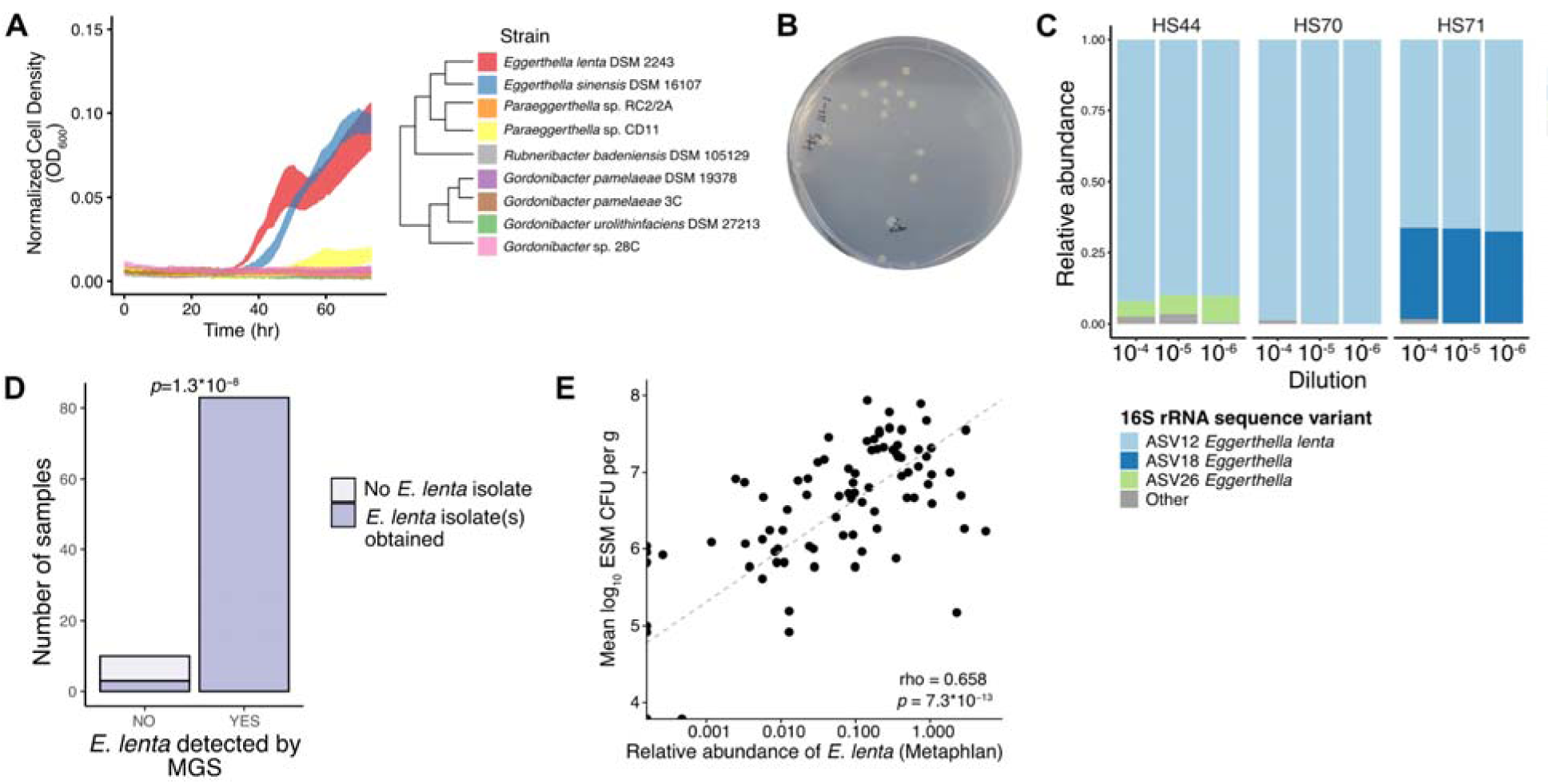
Selective isolation of *E. lenta* from human stool samples. **(A)** ESM supports the anaerobic growth of *E. lenta* and not closely-related members of the *Eggerthellaceae* family (phylogenetic relationships shown from GTDB). **(B)** Example ESM agar plate with *E. lenta* colonies. (**C)** 16S rRNA gene sequencing-based validation of the selectivity of ESM for amplicon sequence variants (ASVs) assigned to genus *Eggerthella* (n=3 stool samples). **(D,E)** Culture-dependent (ESM) and -independent (MetaPhlAn analysis of metagenomic data) detection of *E. lenta* in the ImmunoMicrobiome cohort^31^ show consistent presence/absence **(D)** and abundance **(E)**. Statistics: **(D)** Fisher’s exact test; **(E)** Spearman correlation.

**Figure S2.**
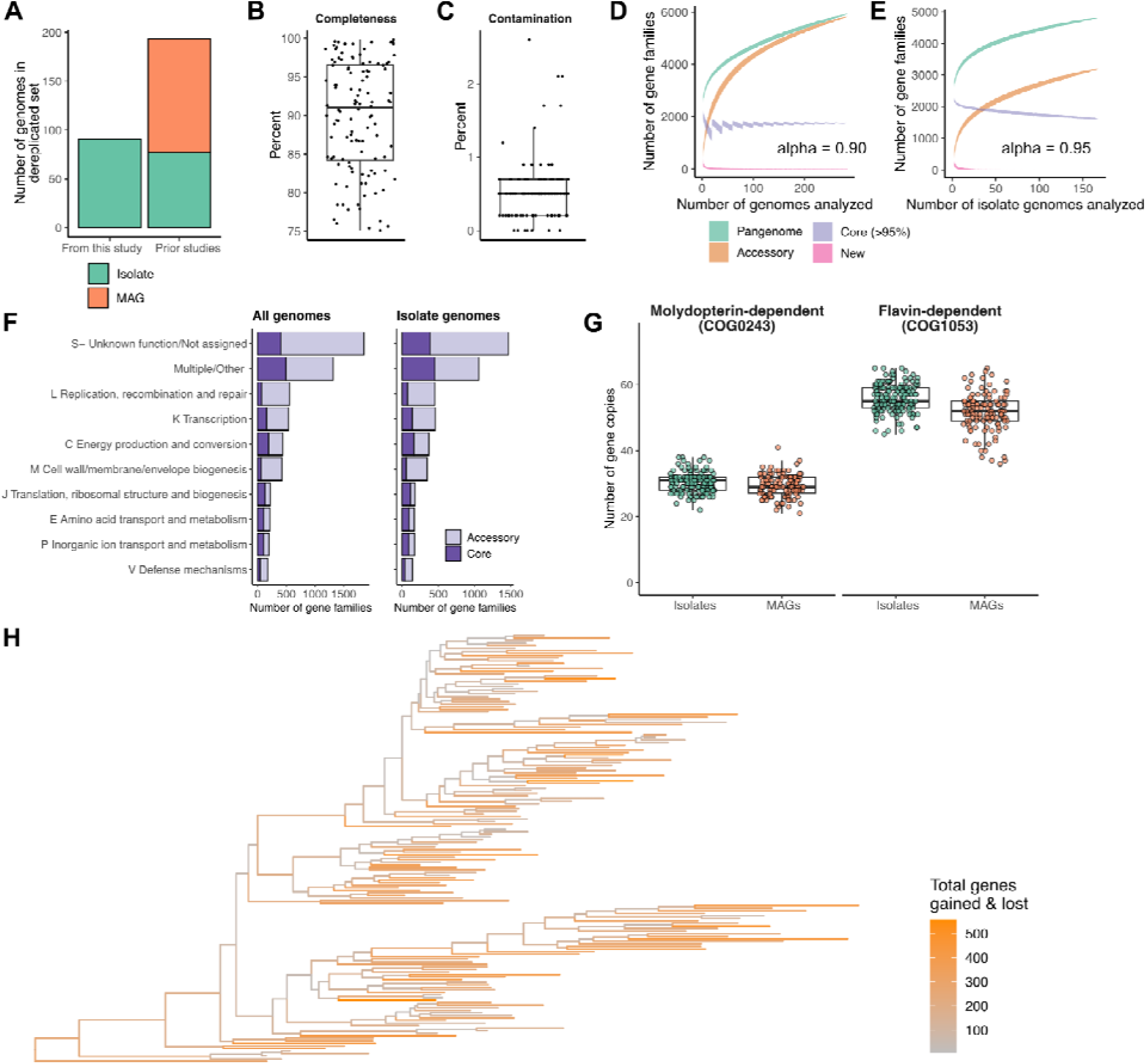
*E. lenta* pangenome variation. **(A-C)** Genomes analyzed in this study, including their sources **(A)** and for MAGs, their completeness **(B)** and contamination **(C)** statistics (BUSCO^79^). **(D,E)** Pangenome sampling curve showing the accumulation of total genes (green), accessory genes (orange), core genes (purple), and new genes (pink) as a function of genomes analyzed for **(D)** all genomes (95% cutoff for core genome inclusion) and **(E)** isolate genomes (100% prevalence cutoff). Band width represents the standard deviation across samplings (*n=*284 for all genomes and 168 for isolates only). Each curve is labeled with the alpha parameter estimated by a Heaps’ law model with 500 iterations^88,89^. Values <1 indicate an open pangenome. **(F)** Distribution of core and accessory gene families (eggNOG) across the most common COG categories for all genomes (95% cutoff for core genome inclusion) and isolate genomes (100% prevalence cutoff). Category S (Unknown function) was combined with unassigned gene families. **(G)** Variation in copy number of genes from two anaerobic reductase superfamilies among isolate genomes and MAGs. **(H)** Maximum likelihood phylogenetic tree (Figure 1A), recolored by the estimated number of gene families gained and lost at each node of the tree (*Panstripe*^90^ phylogenetic model).

**Figure S3.**
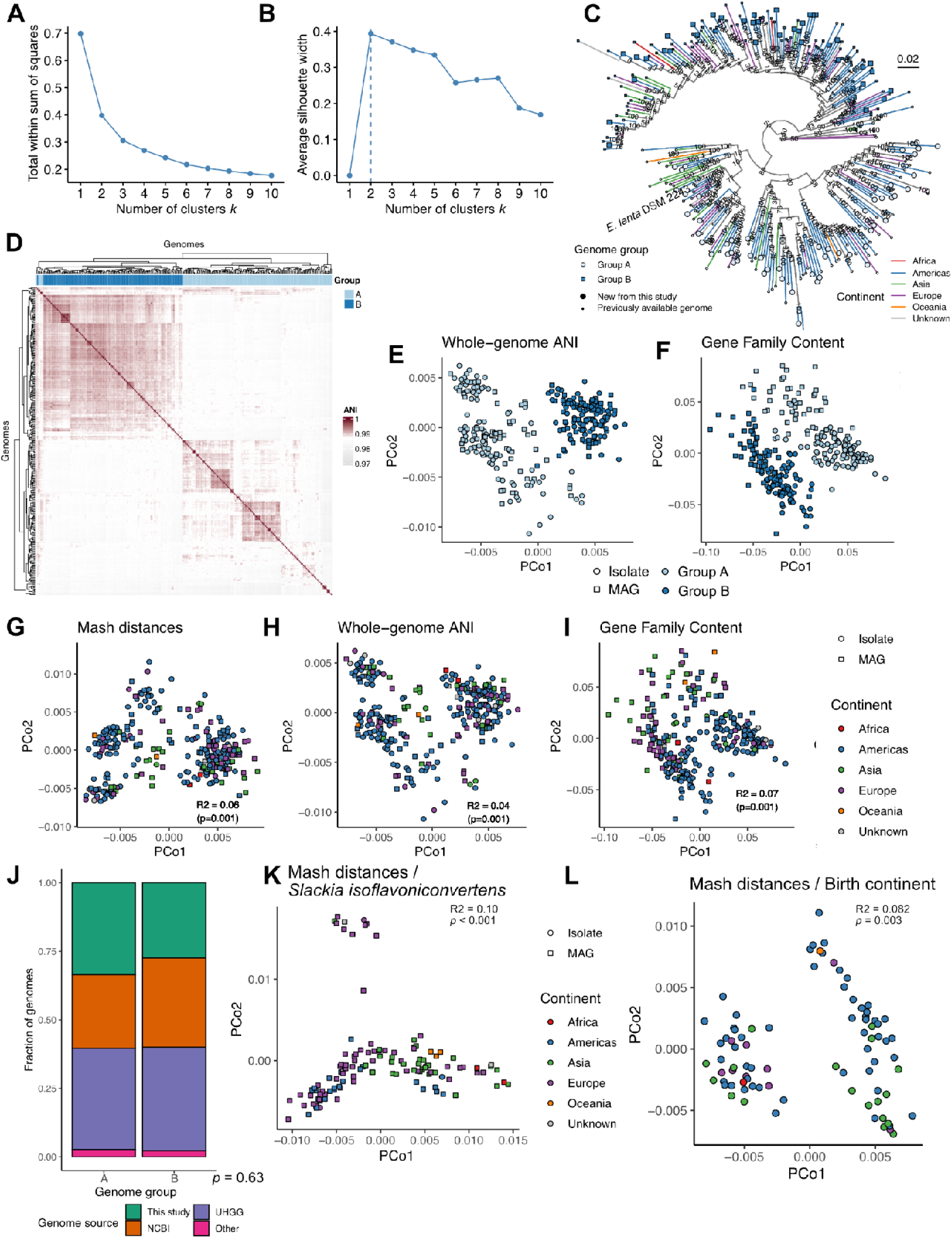
*E. lenta* genomes cluster into two groups with a limited phylogeographic signal. **(A-B)** Elbow (**A**) and silhouette (**B**) plots evaluating the optimal number of genome clusters in our dataset, based on Mash sequence dissimilarity. **(C)** Maximum likelihood phylogenetic tree of the *E. lenta* species based on a core gene alignment generated by *Panaroo*^95^. Bootstrap values were estimated for each node using RAxML. Tips shown in a larger size represent new genomes from this study, while tip shape indicates genome group. Branch colors indicate geographic origin. **(D)** Hierarchical clustering of genomes based on whole-genome average nucleotide identity (ANI), showing a division into two main groups. **(E-F)** Principal coordinates analysis of genomes based on whole-genome ANI (**E**) and eggNOG gene family presence-absence (**F**), illustrating division into groups A and B. **(G-I)** Principal coordinates analysis based on sequence Mash distances (**G**), whole genome ANI (**H**), and eggNOG gene family presence-absence (**I**), as in Figure 1B and **Figure S3E-F**, but colored by geographic origin. Effect sizes and *p*-values are shown for the association between dissimilarity and continent of origin. **(J)** Bar graph showing the fraction of genomes in groups A and B from each of the major sources for this study. *p-*value based on Fisher’s exact test. **(K)** Principal coordinates analysis based on Mash distances of genomes from a related *Eggerthellaceae* species, *Slackia isoflavoniconvertens*. **(L)** Principal coordinate analysis of *E. lenta* genomes from the ImmunoMicrobiome cohort^31^ based on sequence Mash distances, colored by host continent of birth. Effect size and *p-*values in **E-I, K,L** are based on PERMANOVA. Color and shape legends are shared between panels **K** and **L.**

**Figure S4.**
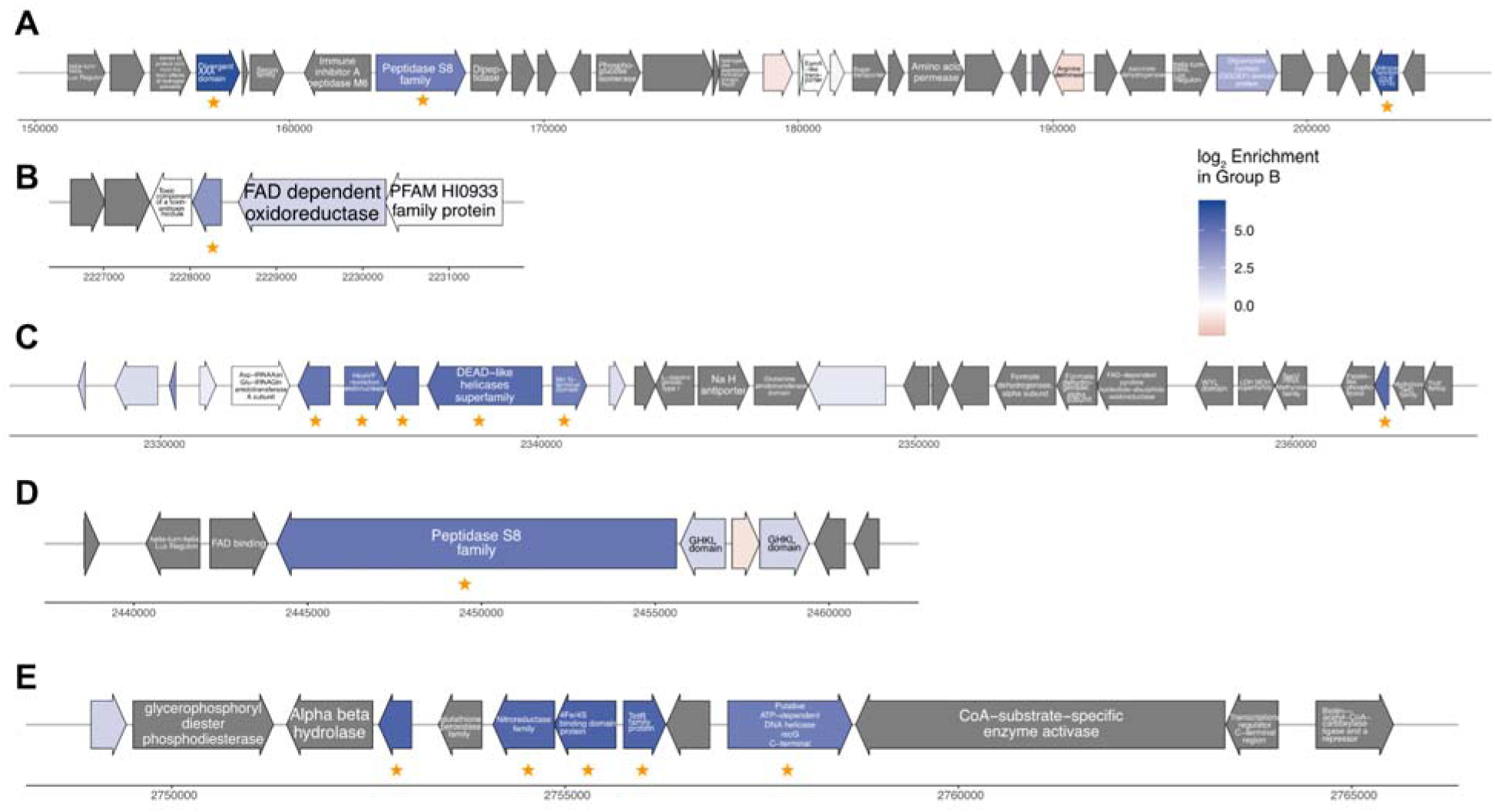
Identification of multiple genomic loci containing Group B enriched genes. **(A-E)** Genomic context for the 16 gene families strongly linked to *E. lenta* Group B (**Table S3, Methods**). Gene maps are based on *E. lenta* APC055-928 and ordered based on coordinates in this genome. Gold stars indicate top Group B-linked genes. Genes are labeled with eggNOG gene family descriptions and colored based on their enrichment in Group B vs. Group A. Genes shown in gray are part of the core genome.

**Figure S5.**
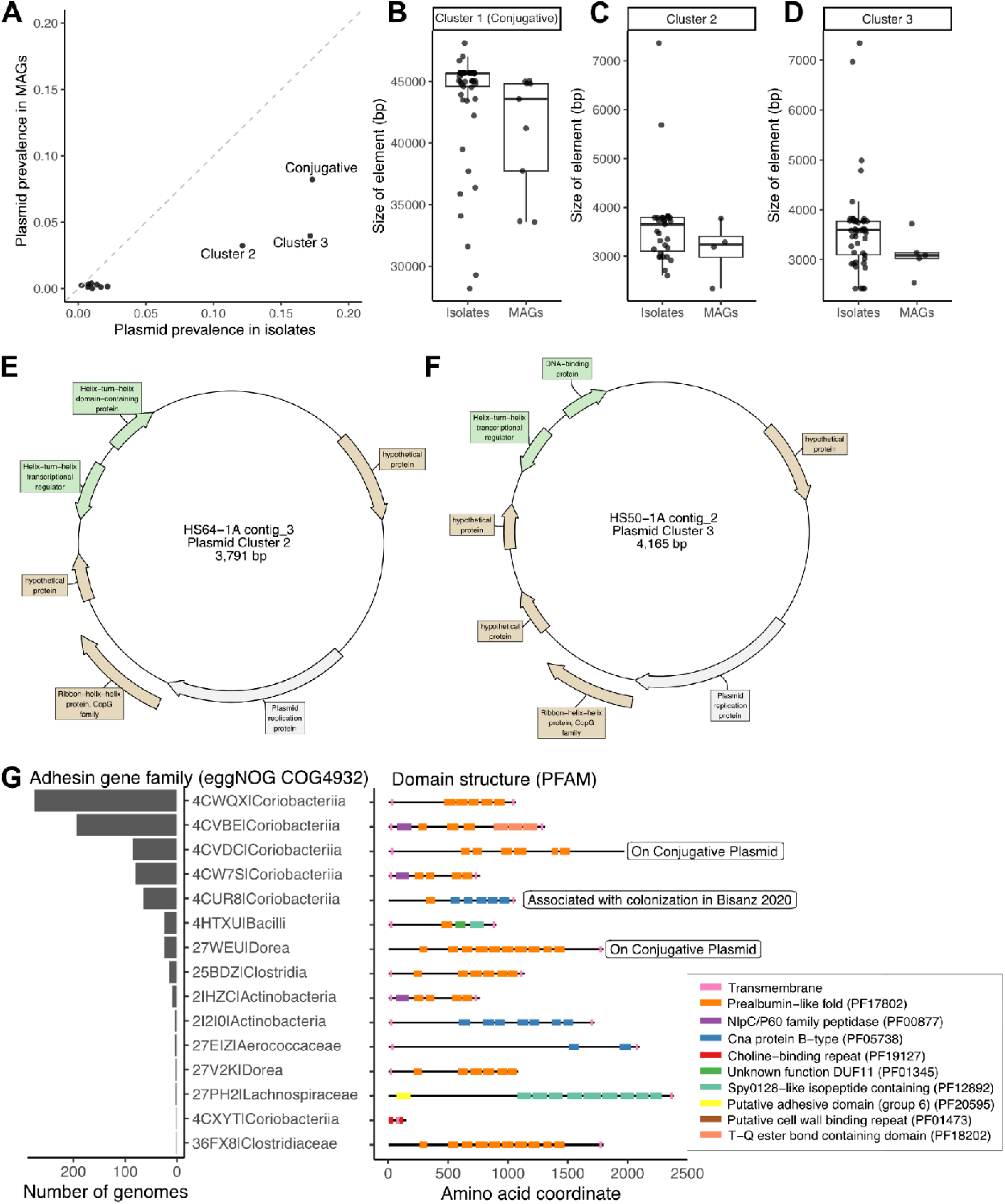
Discovery of three types of plasmids that are widespread in the *E. lenta* species. **(A)** Prevalence of plasmid clusters (sets of closely related plasmids found across genomes, **Methods, Table S5**) across both isolates and MAGs. The three clusters detected in at least 10 genomes are labeled. All clusters had higher prevalence in isolate genomes than in MAGs (below the dashed diagonal line indicating equal prevalence), as expected. **(B-D)** Plasmid element sizes are similar between MAGs and isolates. Boxplots show the distribution of contig sizes among elements in the 3 most prevalent clusters. **(E-F)** Representative gene maps showing gene annotations for the most common small plasmid clusters. Genes are colored by functional category (green = transcription, gray = replication, brown = unknown function). **(G)** Summary of annotated host-binding adhesin gene families found in the *E. lenta* pan-genome (belonging to COG superfamily COG4932). Gene families are labeled with the eggNOG subfamily ID and the associated eggNOG taxonomic assignment. For each gene family, the left panel shows their prevalence in the genome collection, while the right panel shows their PFAM domain structure as annotated by InterProScan (right panel). Two gene families that are only found on the conjugative plasmid are labeled, as is a related gene family that was associated with impaired fitness within the gastrointestinal tract^29^.

**Figure S6.**
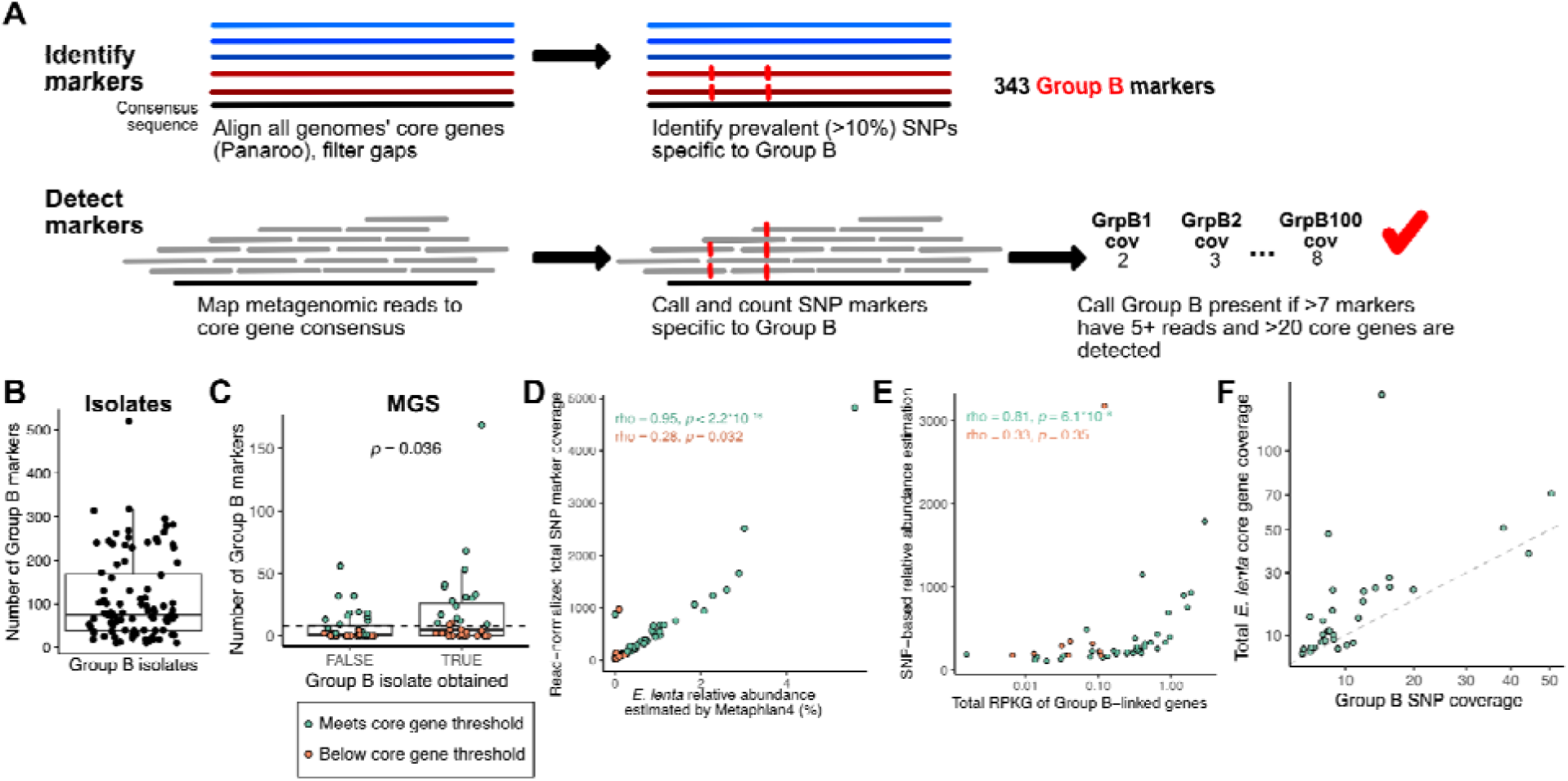
Use of core gene alignment to distinguish *E. lenta* Group B in metagenomic datasets. **(A)** Method used to quantify *E. lenta* Group B. Core genes were retrieved from isolate genomes and aligned to identify SNP markers found in >10% of Group B genomes and absent in the rest of the *E. lenta* species. Metagenomic reads were then aligned to the core gene consensus and the presence of Group B-specific SNP markers was used to determine its presence. **(B,C)** The number of distinct Group B SNP markers varies between isolate genomes in this group **(B)** and is higher in metagenomic data from samples where a Group B *E. lenta* isolate was obtained **(C)**. **(D)** Coverage of all SNP markers is correlated with Metaphlan-based *E. lenta* abundance (Spearman correlation). **(E)** Normalized SNP-based relative abundance of Group B (average coverage of core gene SNP sites per billion reads) is correlated with HUMAnN-based abundance of Group B-linked genes (reads per kilobase per genome equivalent, RPKG). **(C-E)** Colors indicate samples above (green) or below (orange) our minimum detection threshold (>20 core *E. lenta* genes). **(F)** Coverage of Group B SNPs is often lower than total coverage of *E. lenta* core genes. Each point shown represents a sample that met coverage thresholds for Group B detection. The dotted line is *y=x*. Both axes are on a square root scale to visualize low-abundance points.

**Figure S7.**
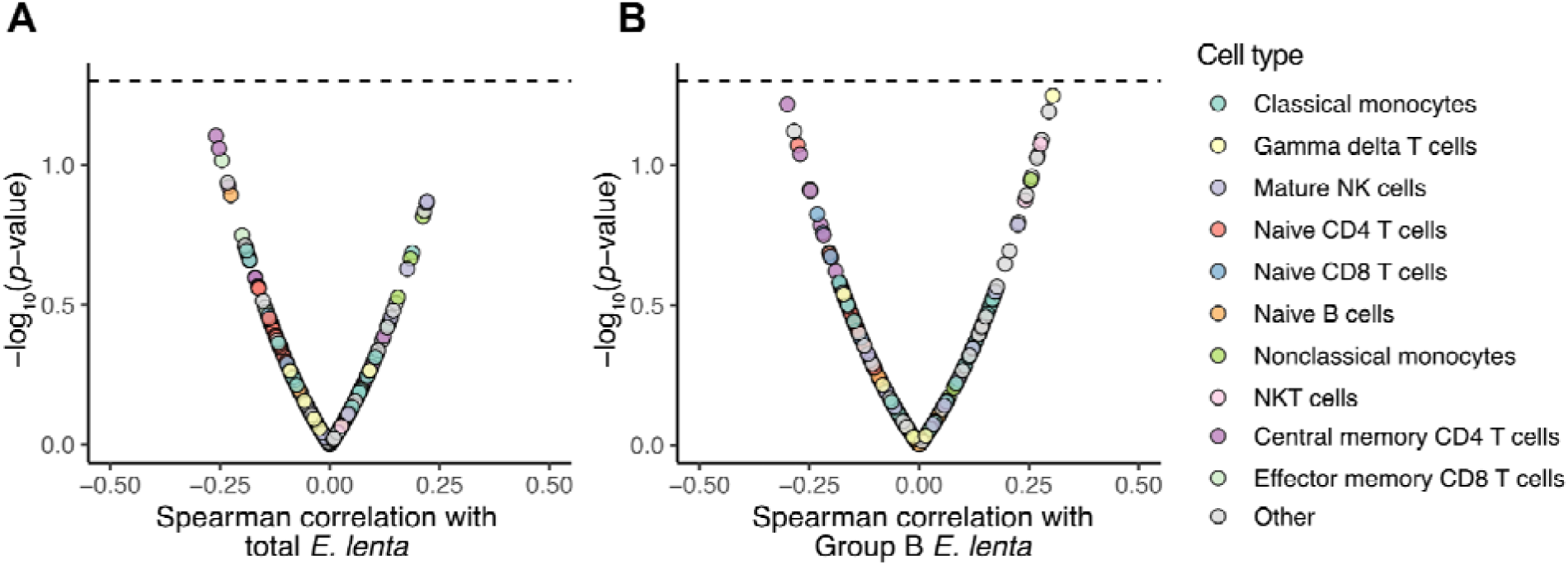
*E. lenta* is not associated with circulating immune cell populations in the ImmunoMicrobiome cohort. Volcano plots showing association of CyTOF cell population scaled cluster frequencies from serum samples, as described^31^, with the SNP-based abundance (average coverage per billion reads) of the *E. lenta* species **(A)** or *E. lenta* Group B (**B**), based on core gene markers. Dotted line represents an uncorrected *p*-value of 0.05.

## SUPPLEMENTAL TABLES

**Table S1. *Eggerthella* selective media (ESM) recipe (1 liter).**

**Table S2. Genome sequencing statistics for isolates newly sequenced in this study.**

**Table S3. 1,425 genomes in the analysis collection (284 *E. lenta*), genome coverage/assembly statistics and metadata for the full collection.**

**Table S4. Prevalent gene families in the *E. lenta* pangenome.**

**Table S5. Gene annotations on the detected *E. lenta* plasmids.**

